# RVQ-Alpha: Bridging Single-Cell Transcriptomics and Large Language Models via Hierarchical Discrete Tokenization and Fact-Aware Reinforcement Learning

**DOI:** 10.64898/2026.04.20.719773

**Authors:** Guangpeng Li, Yue You, Yunlin Fu, Weige Zhou, Feng Tang, Jiaming Kong, Luyi Tian

## Abstract

Single-cell RNA sequencing yields continuous expression profiles, whereas large language models operate over discrete autoregressive sequences, leaving no shared computational interface for language-model reasoning over cell states. Existing approaches either keep cellular information outside the LLM vocabulary, consume context per listed gene, or learn reconstruction codes without gene-level grounding. We introduce **RVQ-Alpha**, which systematically adapts four stages of LLM training (tokenization, supervised fine-tuning, reinforcement learning, and distillation) to single-cell analysis. Multi-codebook **Residual Vector Quantization (RVQ)** lexicalizes each profile into a compact, hierarchical cellular alphabet in the model’s native token stream, while a paired decoder reconstructs the corresponding expression profile. **Evidence-First** supervision grounds these symbols in named genes and expression-linked evidence; **Fact-Aware RLVR** then penalizes contradictory claims to support auditable reasoning. Task-specific RLVR yields strong experts, but a single Mixed policy underperforms them across all four task families. To recover specialist competence in a unified model, we adopt **Multi-Teacher On-Policy Distillation (MOPD)**, which consolidates these experts into one All-in-One checkpoint without retaining a separate policy for each task. On the **CAPSTONE** benchmark, a four-task suite with explicit biological shifts and deterministic ontology-aware graders, the unified checkpoint improves over Mixed on all 12 metrics and remains within 0.005 of the task-routed specialist reference on every primary metric. Overall, RVQ-Alpha provides a unified pipeline for grounded, auditable multi-task single-cell analysis.

## 1 Introduction

Building computational models that can understand and ultimately simulate cellular behavior—the ambition of the *AI Virtual Cell* [1]—is a central goal of computational biology. A necessary foundation for this broader vision is the ability to interpret cellular state from molecular evidence: to identify what a cell is, where it comes from, and how its state changes across health and disease. Single-cell RNA sequencing (scRNA-seq) [2] provides this evidence at cellular resolution, while large language models (LLMs) provide a general interface for integrating context and expressing structured inferences [3, 4]. Connecting them would allow one autoregressive model to reason across both individual cells and multi-cell populations, using the same representational substrate that it uses for language.

The difficulty lies in the representational boundary between these two systems. Gene expression is high-dimensional and continuous, whereas an autoregressive LLM operates over a finite symbolic vocabulary. An effective interface must therefore preserve task-relevant expression structure, remain compact enough for population-level contexts, participate in the model’s native token computation, and retain a route from biological conclusions back to the expression evidence that supports them. These requirements pull in different directions.

Modern single-cell foundation models learn increasingly expressive **continuous representations** from large transcriptomic corpora [5, 6, 7, 8]. Their embeddings provide effective substrates for downstream prediction, although their performance remains task-dependent in systematic comparisons [9, 10, 11]. Because these representations typically terminate in latent states or task-specific heads, they do not by themselves provide a vocabulary-native, reversible interface through which an LLM can read cellular evidence and map the resulting symbols back to an expression frame.

**Text-based interfaces** take a complementary route by serializing expression as ranked gene lists or biological descriptions that pretrained LLMs can read directly [12, 13, 14, 15]. This representation preserves the legibility of named genes, but it collapses expression magnitude into rank order and requires additional context for every feature made explicit—a cost that compounds when many cells must be considered together [16]. Reasoning-oriented systems enrich this textual interface with chain-of-thought supervision or reinforcement learning [17, 18]; connecting the resulting claims to the instance-specific expression profile remains a separate design problem.

**Discrete tokenization** offers a third possibility: represent a cell with a short sequence of symbols that participates directly in autoregressive computation. CellTok [19], for example, places vector-quantized cell codes in an LLM vocabulary and demonstrates the promise of early fusion for population-level analysis. Yet reconstruction-trained codes acquire no explicit gene-level meaning merely by becoming discrete, and a single shared codebook does not explicitly factor representational capacity across successive residual refinements. Discretization makes cellular state writable, but not yet biologically legible.

These interfaces distribute a common burden among compactness, lexical compatibility, and evidential traceability. Our central premise is that a useful cellular alphabet depends on jointly designing its representation and its downstream learning contract. The problem therefore extends beyond constructing a quantizer: the new symbols must be grounded in biological evidence, the claims derived from them must remain inspectable, and task-specific competence must be consolidated without fragmenting deployment across specialist checkpoints.

We introduce **RVQ-Alpha**, which couples residual tokenization, evidence-first semantic grounding, fact-aware verifiable optimization, and specialist-to-generalist consolidation. Figure 1 illustrates its representation and alignment backbone. A *Residual Vector Quantization* (RVQ) tokenizer lexicalizes each cell as eight symbols from eight residual codebooks of 32 entries each (8 × 32 = 256 vocabulary additions). Successive codes refine the representation at a fixed sequence length, while a paired decoder maps the sequence back to the reference expression frame. The resulting bidirectional interface lets the LLM read cellular and textual evidence in one token stream.

**Figure 1.**
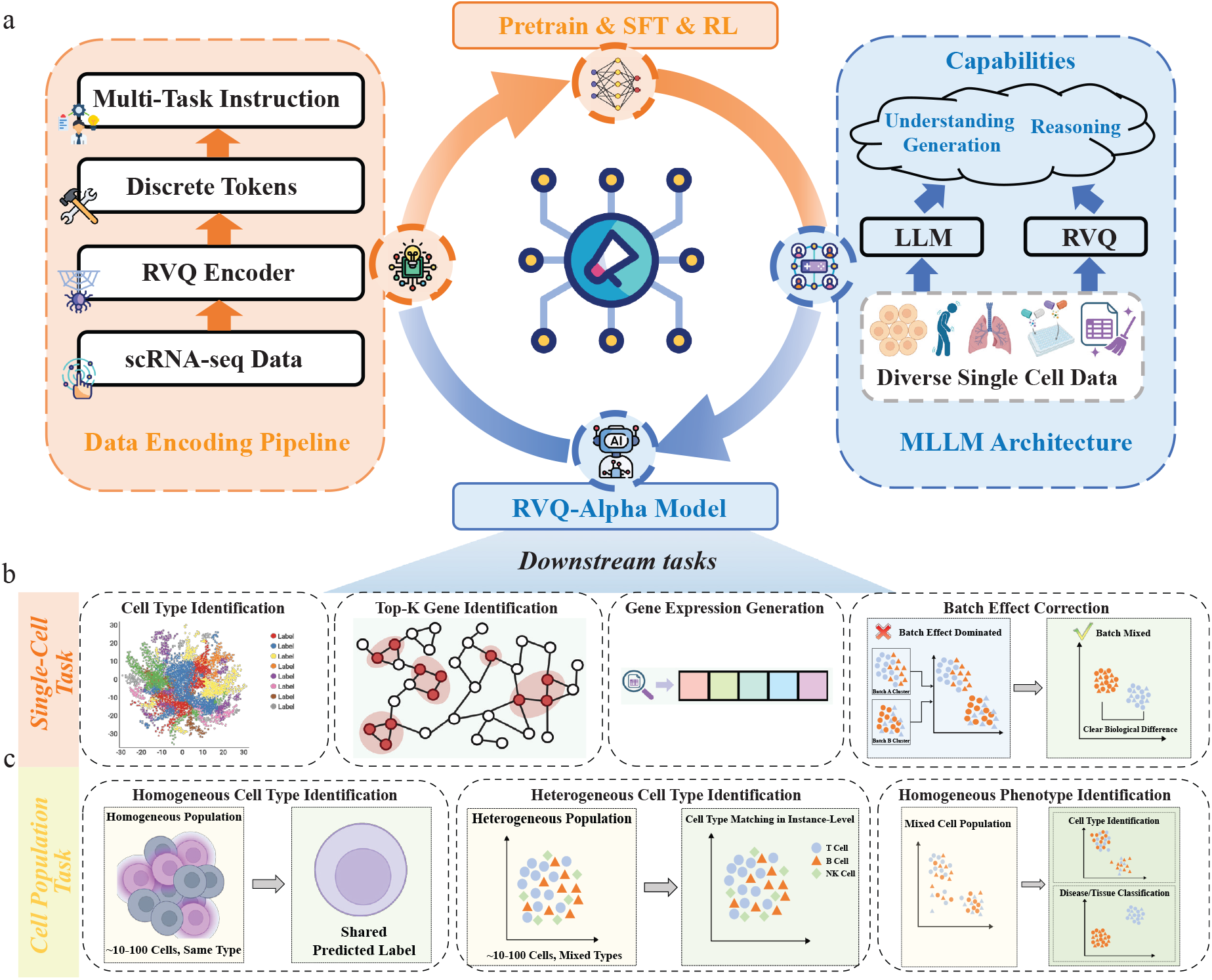
Overview of RVQ-Alpha’s representation and alignment backbone. Top: An RVQ encoder maps scRNA-seq profiles to discrete cellular tokens, which are integrated with an LLM through staged pretraining, supervised fine-tuning, and reinforcement learning. Bottom: Representative single-cell and population-level settings motivate a common interface for reasoning across individual and multicell contexts

Lexicalization establishes compatibility, but newly introduced RVQ symbols begin without the biological semantics required for interpretation. **scCoT-Synth** constructs evidence-before-conclusion reasoning trajectories, and **Feature-Injection SFT** presents the top-*K* expressed genes alongside the RVQ sequence during training. The supervision first exposes the molecular evidence and then derives the biological conclusion, allowing next-token prediction to serve as an implicit cross-modal alignment signal. The resulting tokens serve both as reconstruction indices and as elements of a predictive grammar connected to named genes and structured biological reasoning.

Semantic grounding alone does not ensure that a later rationale remains faithful to the observed cell. We therefore develop **Fact-Aware RLVR** [20], which combines an ontology-aware task reward, structured reasoning assessment, and a target-isolated fact check. The verifier decomposes each rationale into atomic biological claims and checks them against the cell’s expression profile, turning evidence consistency into a fine-grained training signal. Answer quality and evidential support thus become related but distinguishable learning objectives, extending reinforcement learning for biological reasoning [21] from outcome assessment to instance-specific verification.

Verifiable optimization creates strong task specialists, but specialization and deployment impose different demands. Independent policies can adapt to heterogeneous task rewards, whereas direct mixed-task optimization must reconcile those rewards within one policy; retaining every specialist instead requires checkpoint routing at inference time. We separate capability acquisition from capability consolidation through **Multi-Teacher On-Policy Distillation** (**MOPD**) [22, 23]. After specialists are optimized independently, a unified student receives task-routed teacher feedback on trajectories sampled from its own evolving policy, folding complementary capabilities into a single deployable checkpoint.

A shared cellular interface also requires a shared evaluation contract. We therefore establish **CAPSTONE** (*Cell Annotation and Population-level tasks under Shift: a Testbed with Ontology-Normalized Evaluation*), a four-task suite spanning cell-type annotation, tissue and disease-state identification, and population composition analysis across 13 canonical held-out datasets under explicit biological shifts (§4.3). Deterministic ontology-aware graders place every system under the same task-level scoring contract. Within this contract, MOPD improves over mixed-task training on all 12 metrics and remains within 0.005 of the task-routed specialist reference on every primary metric. External comparisons remain task-dependent: MOPD leads single-checkpoint systems on cell-type and tissue macro-F1 and on population-composition detection AUROC, whereas Cell-o1 leads disease-state discrimination and DeepSeek-V4-Pro leads strict composition recovery.

Our main contributions are as follows:

- We introduce a **residual-quantized cellular alphabet** that represents each cell with eight shared-vocabulary symbols and preserves a decoder path back to the reference expression frame (§3.2).
- We develop an **evidence-grounded learning stack**: scCoT-Synth and Feature-Injection SFT connect cellular tokens to named genes through evidence-first trajectories, while Fact-Aware RLVR couples ontology-aware task reward to a target-isolated atomic-claim check (§3.3, §3.4.2).
- We identify a **mixed-versus-specialist gap** and address it with **MOPD**, a separate-then-unify protocol whose consolidated student improves over mixed-task training on all 12 metrics while retaining at least 99.5% of the specialists’ macro-F1 and AUPRC in one checkpoint (§3.5, Table 5).
- We establish **CAPSTONE**, a four-task, expression-traceable suite with explicit biological shifts, deterministic ontology-aware graders, and shortcut controls (§4.3).

**Table 1:** Construction-time verification stack. Layers 1–5 assess the structure and semantic quality of synthesized trajectories; Layers 6–7 enforce representation and nomenclature validity across construction routes. ^†^ Non-blocking quality signal recorded with the CoT sample. ^‡^ Blocking criterion for continued-pretraining samples; failures are regenerated.

| L | Name | Method | Role |
| --- | --- | --- | --- |
| 1 | Context constraint | System-prompt rule | Scope signal<sup>†</sup> |
| 2 | Evidence-first structure | Position check | Soft<sup>†</sup> / Hard<sup>‡</sup> |
| 3 | Forbidden reference | Phrase filter | Soft<sup>†</sup> / Hard<sup>‡</sup> |
| 4 | Reasoning–conclusion | Label extraction | Consistency signal<sup>†</sup> |
| 5 | Semantic rubric | LLM-based multidimensional score | Quality score |
| 6 | RVQ code arithmetic | Offset and range validation | Admission gate |
| 7 | Nomenclature normalization | Curated mapping rules | Admission gate |

**Table 2:**
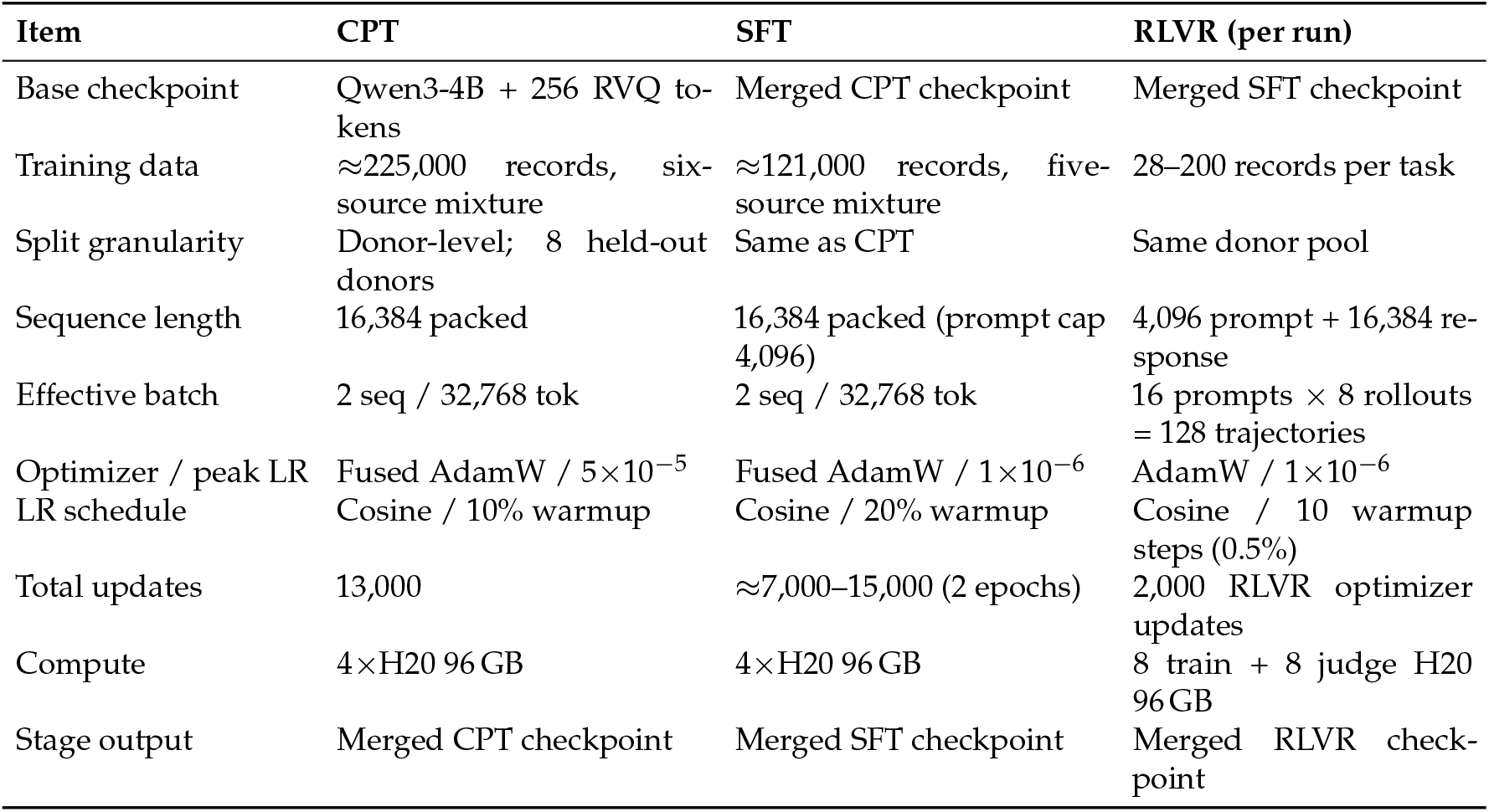
Training protocol before specialist consolidation. Stage-wise settings for CPT, Evidence-First SFT, and each RLVR run. These stages produce the shared initialization and task-specialized policies subsequently used by MOPD.

**Table 3:** Corpus summary: 55-file indexed subset of ScBank. Label counts reflect the harmonized metadata space of the indexed corpus, not the effective class counts of any downstream benchmark.^†^

| Property | Value |
| --- | --- |
| Indexed files | 55 |
| Total cells | 31,481,661 |
| Source study families | ≈45 distinct upstream studies |
| Standardized tissues | 101 |
| Standardized disease states | 103 |
| Standardized cell types | 859 |
| Catalog traceability | 53/55 IDs linked to upstream catalog entries |

<sup>†</sup>Core metadata fields (donor ID, tissue) are present in all 55 files; disease and cell type in 54; sex in 50; age in 42.

**Table 4:**
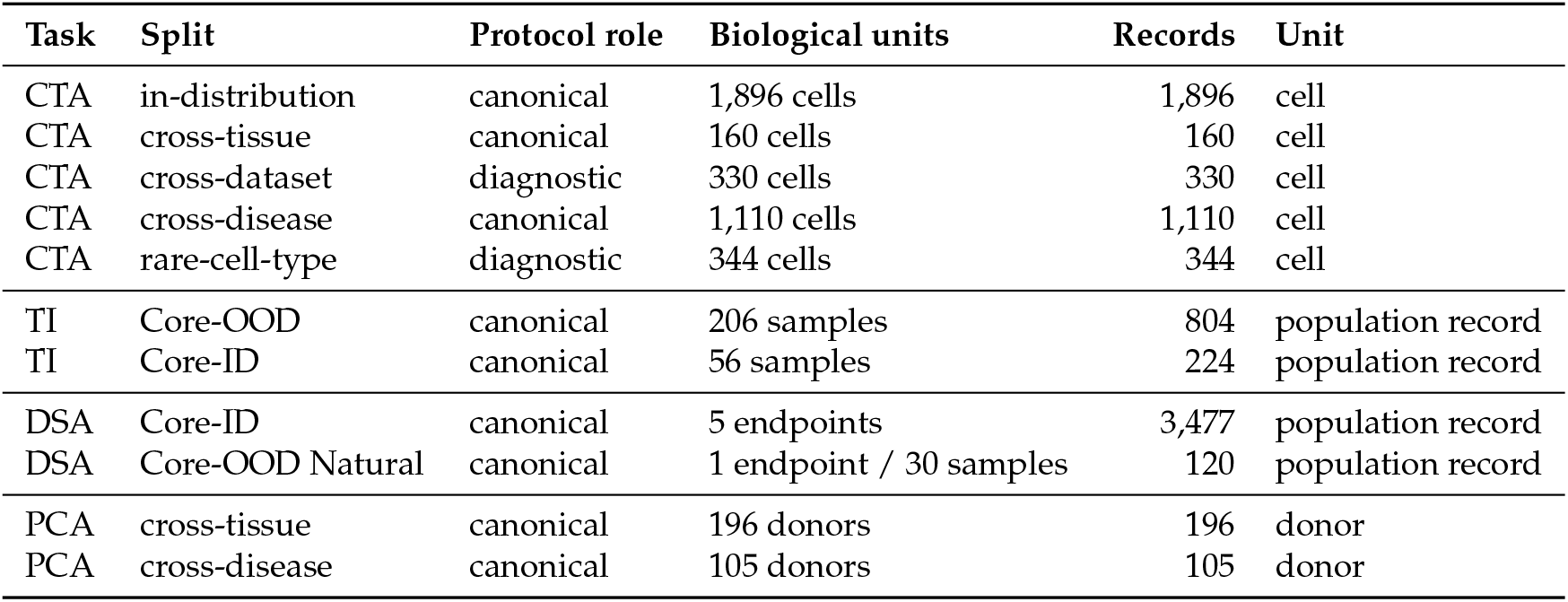
CAPSTONE task and shift inventory. *Biological units* counts independent cells, samples, or donors; *Records* counts reported evaluation instances after nested-*N* expansion. TI and DSA may include several population views of one biological unit, which are not independent biological replicates. Diagnostic splits contribute no record to Table 5.

**Table 5:** Cross-model capability profiles and checkpoint consolidation on CAPSTONE. Headline views are CTA across disease, TI on held-out studies, DSA on natural bags from a held-out study, and an equal-weight macro over the two PCA shifts. *Exact*/*CLDist* denote exact Cell Ontology accuracy/*is - a* distance, and *USim* is UBERON semantic similarity [51]. Strict PCA metrics retain malformed and missing outputs under the worst-case rule. Bold marks the best column value; “—” marks unsupported native workflows. SFT, Mixed-Task, and MOPD are single checkpoints; ^§^ is the task-routed specialist reference, ^‡^ the Qwen3-4B text-input control, and ^†^ the CTA-only C2S-Scale-Gemma-2-2B.

| Model | CTA |  |  | TI |  |  | DSA |  |  | PCA |  |  |
| --- | --- | --- | --- | --- | --- | --- | --- | --- | --- | --- | --- | --- |
|  | mF1↑ | Exact↑ | CLDist↓ | mF1↑ | BAcc↑ | USim↑ | AUPRC↑ | AUROC↑ | BAcc↑ | sJSD↓ | DetAUC↑ | sDom↑ |
| *General LLMs* |  |  |  |  |  |  |  |  |  |  |  |  |
| Qwen3-4B-Thinking-2507<sup>‡</sup> | 0.069 | 0.050 | 3.904 | 0.236 | 0.209 | 0.700 | 0.306 | 0.437 | 0.483 | 0.621 | 0.559 | 0.333 |
| Qwen3-30B-A3B-Thinking-2507 | 0.232 | 0.241 | 2.807 | 0.417 | 0.333 | 0.919 | 0.319 | 0.462 | 0.494 | 0.092 | 0.670 | 0.878 |
| Qwen3.6-35B-A3B | 0.257 | 0.264 | 2.749 | 0.452 | 0.369 | 0.890 | 0.336 | 0.508 | 0.505 | 0.169 | 0.700 | 0.840 |
| Kimi-K2 | 0.210 | 0.250 | 2.462 | 0.267 | 0.206 | 0.858 | 0.489 | 0.708 | 0.675 | 0.188 | 0.630 | 0.788 |
| GPT-OSS-20B | 0.207 | 0.147 | 2.560 | 0.346 | 0.288 | 0.262 | 0.324 | 0.478 | 0.489 | 0.596 | 0.571 | 0.405 |
| GPT-OSS-120B | 0.203 | 0.165 | 2.496 | 0.435 | 0.364 | 0.695 | 0.331 | 0.494 | 0.498 | 0.669 | 0.579 | 0.327 |
| Doubao-Seed-2.0-Pro | 0.261 | 0.231 | 2.095 | 0.569 | 0.445 | 0.884 | 0.350 | 0.541 | 0.522 | 0.046 | 0.745 | 0.973 |
| DeepSeek-V4-Flash | 0.325 | 0.275 | 2.046 | 0.668 | 0.602 | 0.834 | 0.347 | 0.530 | 0.512 | 0.058 | 0.755 | 0.968 |
| DeepSeek-V4-Pro | 0.295 | 0.231 | 2.196 | 0.432 | 0.379 | 0.810 | 0.321 | 0.471 | 0.496 | **0.045** | 0.769 | **0.976** |
| *Domain models* |  |  |  |  |  |  |  |  |  |  |  |  |
| Cell-o1 | 0.197 | 0.223 | 2.515 | 0.218 | 0.196 | 0.728 | **0.615** | **0.810** | 0.507 | 0.812 | 0.536 | 0.198 |
| CellTok | 0.261 | 0.374 | 2.308 | — | — | — | 0.370 | 0.550 | 0.518 | 0.429 | 0.521 | 0.536 |
| C2S-Scale-Gemma-2-2B<sup>†</sup> | 0.237 | 0.170 | 2.773 | — | — | — | — | — | — | — | — | — |
| CellWhisperer | 0.078 | 0.042 | 4.520 | 0.242 | 0.165 | 0.752 | 0.317 | 0.430 | 0.425 | 0.075 | 0.725 | 0.938 |
| *Ours* |  |  |  |  |  |  |  |  |  |  |  |  |
| RVQ-Alpha SFT | 0.281 | 0.323 | 2.192 | 0.569 | 0.505 | 0.874 | 0.491 | 0.674 | 0.589 | 0.342 | 0.718 | 0.903 |
| RVQ-Alpha Mixed-Task | 0.309 | 0.355 | 2.086 | 0.634 | 0.566 | 0.906 | 0.553 | 0.741 | 0.643 | 0.287 | 0.752 | 0.948 |
| RVQ-Alpha Specialist<sup>§</sup> | **0.329** | **0.378** | **2.021** | **0.676** | **0.609** | **0.930** | 0.607 | 0.800 | **0.683** | 0.246 | **0.778** | 0.974 |
| RVQ-Alpha MOPD | 0.328 | 0.377 | 2.030 | 0.674 | 0.607 | 0.927 | 0.604 | 0.796 | 0.681 | 0.251 | 0.775 | 0.972 |

## 2 Related Work

Research on language-enabled single-cell modeling can be read as a sequence of interface decisions: how a cell is represented, how that representation enters a language model, how biological meaning is attached to the resulting symbols, and how the resulting policy is verified and consolidated. We organize the literature along these decisions, moving from expression-native encoders to language interfaces, discrete cellular alphabets, and verifiable post-training.

### 2.1 Expression-Native Single-Cell Foundation Models

Expression-native foundation models established the transcriptome itself as a scalable pretraining substrate. Early systems differed chiefly in how they imposed learnable structure on unordered expression profiles: scGPT [5] uses generative masked-expression pretraining over 33 million cells; Geneformer [6] converts expression into rank-value sequences and learns gene-network structure through attention; and scBERT [24] adapts bidirectional pretraining through expression binning. The same program has since expanded in scale and scope through scFoundation [7] (100M parameters and 50M cells), CellFM [25] (800M parameters and 100M cells), and scPRINT-2 [26] (350M cells spanning 16 organisms).

Scaling has been accompanied by architectural diversification. CellPLM [27] models cells as tokens and tissues as sentences for spatial integration; GeneMamba [28] replaces quadratic attention with state-space modeling; AIDO.Cell [29] extends dense modeling to the full transcriptome; UCE [8] transfers protein-language embeddings to zero-shot cell mapping; and Nicheformer [30] and SCimilarity [31] target, respectively, joint dissociated–spatial representation and atlas-scale metric retrieval. Systematic evaluations nevertheless complicate a simple scale narrative: performance leadership varies by task, linear baselines can remain competitive, and representation choices become especially consequential under distribution shift [9, 10, 11, 16]. Taken together, this line advances cellular representation learning; its downstream contract is generally a continuous embedding or task-specific head, whereas RVQ-Alpha studies a vocabulary-level cellular alphabet that can inhabit the same autoregressive stream as text.

### 2.2 Language Interfaces for Single-Cell Reasoning

Language-oriented systems approach the same interface problem from the opposite direction: they preserve the capabilities of a pretrained language model and translate expression into a form it can consume. Serialization makes gene identity directly legible to the model: Cell2Sentence [12] renders expression profiles as ranked-gene sentences, an approach subsequently scaled to 27B parameters [13]; CELLama [32] combines ranked genes with metadata; and scMulan [33] represents genes and expression values as prompted entity–value tuples. Continuous interfaces instead align learned cellular representations with language: LangCell [14] and CellHermes [15] use cross-modal pretraining, while InstructCell [34] and CellWhisperer [35] extend the interface toward instruction following and conversational exploration. Stack [36] further combines tabular attention with in-context learning at a scale of 149 million cells.

These designs expose a recurring trade-off. Projection-based interfaces can summarize a cell in a few continuous vectors, but the resulting state remains outside the model’s lexical vocabulary; gene serialization inherits the semantics of existing language tokens, but its context cost grows with every piece of expression evidence exposed. Single-cell language interfaces thus purchase compatibility in two currencies—adapter capacity or sequence length. The remaining challenge is an interface that is simultaneously compact, vocabulary-native, and traceable to gene-level evidence.

### 2.3 Discrete Lexicalization of Cellular State

Discrete tokenization is more than a compression device: it lexicalizes a continuous modality, allowing its states to participate in the same autoregressive grammar as text. VQ-VAE [37] supplied the foundational quantized-autoencoder construction for images, while neural audio codecs such as SoundStream and EnCodec [38, 39] showed how Residual Vector Quantization can distribute information across a sequence of progressively refined codebooks. Related biological work has also begun to discretize structured molecular objects, including geometric byte-pair encoding for protein structures [40].

Within single-cell analysis, CellTok [19] is the closest antecedent: its VQ-VAE codes enable early-fusion integration with Qwen2.5 and support population-level analysis without first spelling a cell as a long gene list. A shared alphabet resolves interface compatibility, however, not semantic grounding by itself. CellTok uses a single shared codebook optimized for expression reconstruction, whereas RVQ-Alpha makes a residual cascade the interface, so that successive codebooks refine the cellular state and a prefix remains a usable approximation (§5.3). Evidence-First supervision then couples those vocabulary tokens to gene-level evidence, and expression-grounded verification governs how the resulting symbols enter biological claims. In this formulation, discretization begins the interface design rather than ending it: token geometry, biological interpretation, and post-SFT reasoning are treated as connected problems.

### 2.4 Verifiable Post-Training and Capability Consolidation

Once cellular states enter the token stream, the bottleneck shifts from representation to epistemic discipline: biological rationales must be checked against cell-specific evidence, and task-specific gains must survive consolidation into one policy. CellReasoner [17] uses 380 chain-of-thought exemplars to elicit marker-level reasoning for cell-type annotation, Cell-o1 [18] applies reinforcement learning with batch-level rewards, and OmniCellAgent [41] broadens the setting to multi-agent analysis and hypothesis generation. At the methodological boundary, DeepSeek-R1 [20] established the broader RLVR template; rbio1 [21] uses biological world models as soft verifiers; and VCWorld [42] combines structured biological knowledge with interpretable perturbation reasoning. These systems develop complementary forms of supervision—demonstrations, batch rewards, tool orchestration, world-model feedback, and structured knowledge—while RVQ-Alpha asks whether the individual claims in a cellular rationale agree with the expression evidence for the instance being judged.

Verification alone does not resolve the deployment fragmentation created by task specialization. Multi-teacher on-policy distillation in general-purpose models [22] and specialist-to-generalist distillation in medicine [23] instead follow a separate-then-unify principle: specialists retain task-specific competence during optimization, while a student absorbs their feedback along its own evolving trajectories. RVQ-Alpha adapts this principle to task-routed single-cell reasoning, consolidating independently optimized specialists without requiring checkpoint switching at inference time.

Evaluation resources likewise differ in the contract they impose between biological input and model output. SOAR [43] centers marker-list cell-type annotation; SC-Arena [44] evaluates multiple cell-language tasks over ranked-gene sentences; CellVerse [45] emphasizes multi-omics question answering; scBench [46] evaluates agentic scRNA-seq analysis; and VCWorld targets perturbation reasoning. Across these five resources, CAPSTONE addresses an interface-level gap by pairing text and RVQ views of the same biological instance under a deterministic, ontology-aware grading contract (§4.3). Together, these lines expose an integration problem: a cellular representation must be compact enough to inhabit the LLM alphabet, grounded enough to support gene-level justification, trainable under verifiable rewards, and consolidatable across tasks.

## 3 Method

RVQ-Alpha organizes cellular representation and biological reasoning as a sequence of linked learning problems. An upstream RVQ tokenizer first converts an expression profile into a compact cellular alphabet; continued pretraining and Evidence-First supervision then connect that alphabet to named genes and structured biological inference; Fact-Aware RLVR develops task-specialized policies under answer- and evidence-conditioned feedback; and Multi-Teacher On-Policy Distillation (MOPD) consolidates those specialists into one student policy. Figure 1 summarizes this checkpoint lineage.

### 3.1 Problem Formulation and Framework Roadmap

#### Inputs and outputs

Let **x** ∈ ℝ^*G*^ denote a single-cell gene expression profile reconciled to a fixed, positionally aligned reference frame of *G* = 36,601 GRCh38 genes, and let *P* = {**x**_1_, …, **x**_*N*_} denote a population of *N* cells. Each sample is paired with an instruction **t** that specifies the task and its biological context. The language model predicts a response *y*, which may be a categorical label, a ranked gene list, a population-level summary, or a CoT reasoning trace; the paired cellular decoder separately maps an RVQ sequence back to the reference expression frame.

#### Analysis tasks and training routes

The task surface spans four single-cell analyses—cell-type, tissue, and disease-state identification, together with marker-gene ranking—and a population route that combines per-cell annotations into a population-level summary. Each single-cell output pairs its answer with a CoT rationale.

We distinguish the partitions used at successive points in the learning system. Synthesis tasks specify which trajectories scCoT-Synth constructs, SFT routing families determine how supervised records are formatted and sampled, and specialist or teacher routes identify the policies used during RLVR and MOPD. Each count is therefore qualified by its role in data construction, optimization, or evaluation.

#### Checkpoint lineage

The framework has three nested levels of optimization. First, ℒ _RVQ_ trains the upstream tokenizer and decoder while defining the cellular alphabet (§3.2). Second, the LLM passes through continued pretraining, Evidence-First supervised fine-tuning under ℒ_SFT_, and task-specialized RLVR under sequence-level rewards; §3.3 specifies the grounding stage, and Appendix D gives the policy objective. Third, MOPD uses the resulting specialists as frozen teachers and transfers their routed distributions into a unified student (§3.5). Each handoff begins from the checkpoint produced by the preceding phase, while the upstream RVQ encoder remains frozen throughout LLM adaptation and consolidation.

### 3.2 A Vocabulary-Native Cellular Alphabet

A cellular tokenizer for language-model reasoning must satisfy three requirements simultaneously. It must compress each expression profile enough to place tens of cells beside natural-language instructions, preserve variation from broad lineage to fine-grained state, and expose the resulting symbols through the LLM’s native token interface. The third requirement is central to the design: routing cellular information through the shared embedding table lets the ordinary autoregressive objective shape how cellular and textual symbols interact, without inserting a separately learned cross-modal adapter between expression and reasoning.

Text serializations such as Cell2Sentence [12] preserve the legibility of named genes but require 100–500 tokens per cell, while flat VQ-VAE tokenizers such as CellTok [19] compress a cell into 32 assignments from one shared codebook [37]. These alternatives expose the underlying trade-off between lexical transparency, sequence length, and allocation of discrete capacity. RVQ-Alpha addresses it with successive residual refinement and a fixed representation of eight cellular symbols per cell.

#### Residual tokenizer design

A parametric encoder *f*_enc_ maps gene expression to a *D*-dimensional latent **z** = *f*_enc_(**x**). Given *N*_*c*_ codebooks 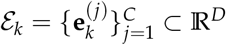, the tokenizer approximates **z** through successive residual assignments:

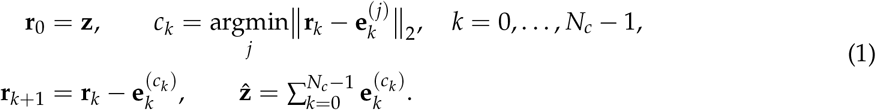

The decoder reconstructs 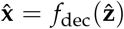 in the same aligned gene space. Tokenizer training combines global reconstruction, reconstruction on coordinates masked from the encoder input, and regularized vector quantization:

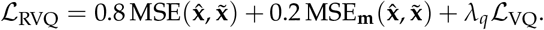

Here 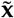 is the reconstruction target and **m** marks coordinates withheld from the encoder. The global term preserves the overall expression profile, whereas the masked-coordinate term requires the latent representation to retain signal that supports recovery from the observed context. The commitment and embedding terms in ℒ_VQ_ couple continuous residuals to their discrete representatives, and its usage regularizer discourages collapse onto a small subset of codes. Supplementary F gives the full architecture, augmentation curriculum, and objective.

We use *N*_*c*_ = 8 codebooks with *C* = 32 entries each, selected by jointly considering reconstruction, code utilization, and label-based clustering on held-out cells (§6.1). The residual recursion gives the codebooks distinct approximation roles: each stage models the residual left by its predecessors, so early assignments capture the largest remaining directions in latent space and later assignments refine progressively smaller residuals. This ordering creates a coarse-to-fine statistical hierarchy. Whether that hierarchy aligns with lineage, cell type, and cellular state is an empirical question, which we examine through codebook-level analyses in §6.

#### Vocabulary-native integration

The *N*_*c*_ × *C* code grid is enumerated into a contiguous block of token identifiers and embedded through the same input–output interface used for text. Each selected pair (*k, c*_*k*_) receives the identifier

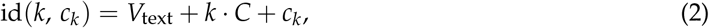

where *V*_text_ is the text-vocabulary offset established when the new symbols are added. This construction introduces 256 cellular symbols and represents each cell with one symbol from each of the eight codebooks; cell- and population-level delimiters mark the boundaries between cells and cell groups. The resulting eight-symbol footprint is one quarter of CellTok’s 32 assignments and substantially shorter than a text-serialized top-gene list, excluding instruction-level tokens shared across cells. The compact interface makes population prompts with tens of cells feasible while preserving a decoder path to the aligned expression frame.

Figure 2 depicts the resulting dual-tokenization interface. Natural language passes through the BPE tokenizer, cellular profiles pass through the RVQ tokenizer, and both streams meet in the LLM’s shared autoregressive computation. This shared interface is the substrate on which the grounding objectives below connect cellular symbols to gene-level language.

**Figure 2.**
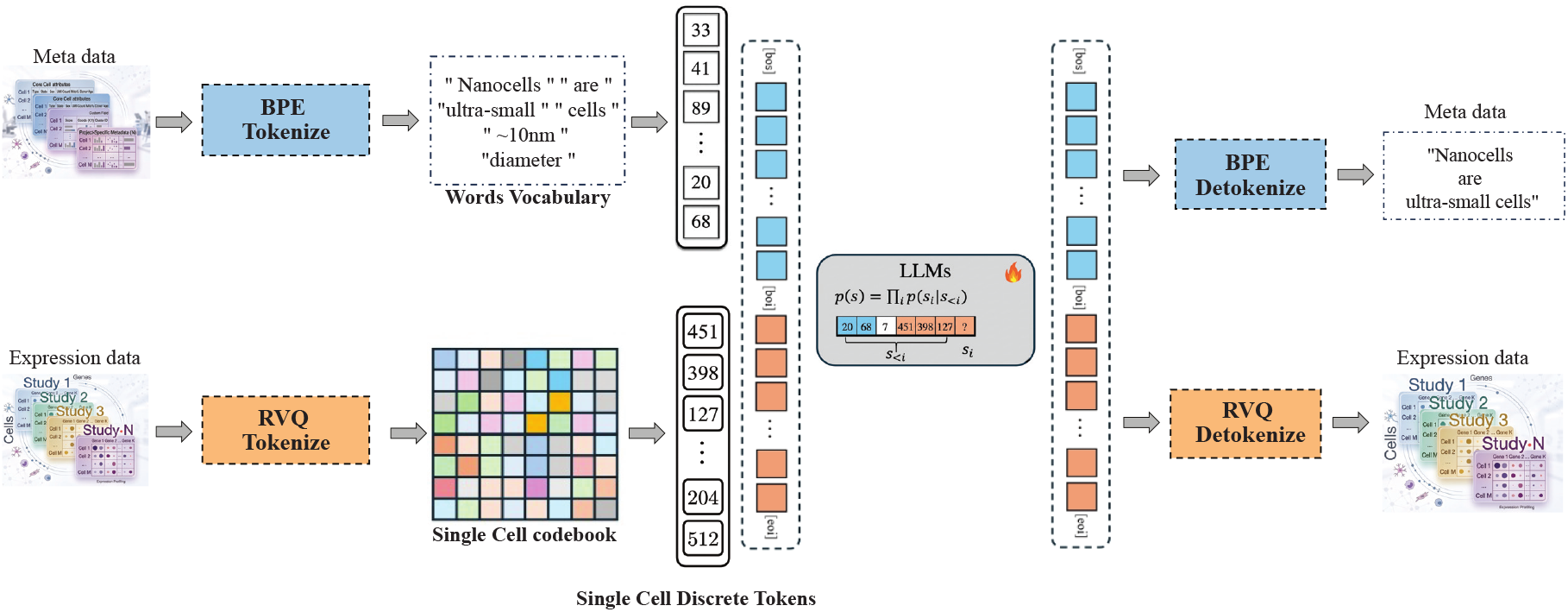
Core dual-tokenization architecture. Text instructions are processed by a BPE tokenizer, while aligned gene-expression profiles are mapped to compact cellular sequences by the RVQ tokenizer. The two token streams enter the same autoregressive LLM, and the paired RVQ decoder retains a path from cellular symbols back to the reference expression frame.

### 3.3 Evidence-First Semantic Grounding

Vocabulary integration makes cellular codes available to the LLM, but the 256 newly introduced symbols begin without the gene-level semantics carried by pretrained language representations. RVQ-Alpha addresses this semantic cold start through two complementary forms of supervision. scCoT-Synth constructs trajectories in which molecular evidence precedes biological inference, while Feature-Injection SFT places top-expressed gene names beside the RVQ sequence during supervised grounding. Together they create a lexical scaffold through which next-token prediction can associate the cellular alphabet with named genes and structured reasoning.

#### Evidence-First learning contract

Every reasoning trajectory follows a directed evidence–inference chain:

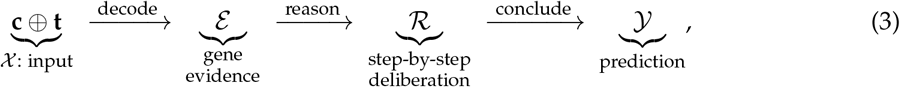

where **c** = (*c*_0_, …, *c*_7_) is the cellular code sequence and **t** is the task instruction. The response first states gene evidence ℰ, then uses that evidence in a reasoning trace ℛ, and finally emits the prediction *Y* Gene-name tokens are therefore supervised at positions that attend to the preceding RVQ sequence, providing a direct learning pathway from lexical gene concepts to the new cellular embeddings.

#### Task-conditioned evidence

scCoT-Synth constructs each trajectory from a task-conditioned *evidence card*. The card may include expression-ranked genes, known markers, transcription-factor activity, functional programs, and competing-lineage counterevidence, but it admits a field only when that field can be obtained without resolving the target of the current task. The same measurement may consequently be legitimate evidence for one question and target-equivalent leakage for another. Constructing the card per task defines the factual material available to the teacher and the boundary against which its claims can be audited.

#### Generator–Evaluator–Refiner loop

A replaceable generator receives **c, t**, and the evidence card and proposes a candidate trajectory. A construction-time evaluator resolves the trajectory into atomic claims and checks whether each claim is supported by the visible card, properly grounded, and scoped to RNA-level evidence. The evaluator returns constraints rather than authoring a replacement, allowing the generator to revise against the same evidential boundary.

Each revision is assessed under the same admission predicate, and the pipeline retains the strongest admissible round rather than defaulting to the final rewrite. Label agreement is excluded from the refinement criterion, so repair operates on evidence quality and reasoning structure instead of recovering the target indirectly. Deterministic checks additionally identify direct label references, malformed evidence ordering, invalid RVQ arithmetic, and noncanonical nomenclature.

The privileged evidence card serves data construction only. Once a trajectory is admitted, the card and its audit trace are removed; the student receives the cellular sequence and task instruction, while the assistant target preserves gene names as claims that must be recovered from the RVQ context. For population trajectories, per-cell evidence establishes the cell-level decisions before aggregation, so the population summary explains those decisions without revising them.

#### Multi-granularity synthesis and construction-time verification

The synthesis engine distributes grounding across cell-level annotation, population-level inference, and extended explanatory contexts. All synthesis routes share the task-specific availability test and role-separated construction loop, while their output schemas determine which structural checks apply. Table 1 distinguishes this offline corpus-admission stack from the reward-time judges and factuality verifier introduced later.

The construction-time semantic evaluator scores evidence grounding, logical rigor, factual accuracy, and uncertainty calibration, using a separate profile for continued-pretraining narratives. These scores rank admissible candidates and characterize corpus quality; they do not become the post-training reward used by RLVR.

#### Feature-Injection SFT

During supervised grounding, Feature-Injection SFT presents the top-*K* expressed gene names alongside the RVQ sequence and task instruction. This co-exposure gives the new cellular symbols an explicit lexical scaffold: the model observes which named molecular features accompany a code sequence while learning to generate evidence before its conclusion. Gene injection is used to establish the alignment during training; at inference time, the model must recover the relevant gene concepts from the cellular symbols and instruction without receiving the injected list.

Continued pretraining first exposes the vocabulary-expanded model to RVQ–text descriptions across multiple linguistic registers, establishing broad associations between cellular codes and biomedical language. Evidence-First SFT then turns those associations into structured task behavior. It updates the LLM from the continued-pretraining checkpoint while keeping the upstream RVQ encoder frozen, and computes loss only on assistant positions:

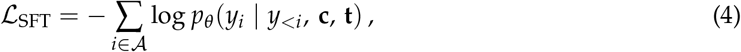

where *A* indexes assistant-token positions, *θ* denotes the LLM parameters, and *y*_*i*_ is the target token at position *i*. Restricting supervision to the assistant response trains the model to reconstruct evidence and formulate conclusions rather than reproduce the prompt.

#### Gradient-mediated alignment

The Evidence-First ordering creates a direct gradient pathway from gene-evidence positions to the cellular embeddings through attention:

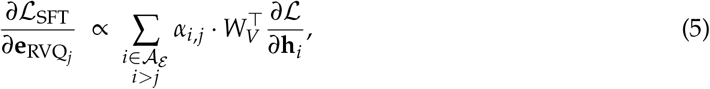

where *A*_ℰ_ indexes gene-evidence token positions and *α*_*i,j*_ is the attention assigned from evidence position *i* to cellular position *j*. Within a layer, stronger attention to a cellular symbol at a gene-prediction position increases the corresponding supervisory signal. This value-path contribution is one of several gradient routes connecting the two modalities, and it explains how ordinary autoregressive training can encourage gene-linked cellular representations without an explicit projector. The resulting checkpoint supplies both the structured responses and the gene-evidence anchors required by Fact-Aware post-training.

### 3.4 Fact-Aware Specialist Post-Training

The grounded SFT checkpoint establishes structured biological response generation. Specialist post-training adds comparison among multiple candidate answers and outcome-dependent correction through a composite sequence reward. Its task component scores answer semantics and reasoning structure, while its factuality component contributes a bounded adjustment for auditable contradictions. We optimize the specialists with Group Sequence Policy Optimization (GSPO) [47], whose response-level importance ratio aligns the policy update with a reward assigned to the complete answer and evidence chain. Appendix D gives the group-normalized advantage, comparison with token-level GRPO, and clipped surrogate; the two reward components are defined below.

#### 3.4.1 Ontology-Grounded Task Reward

Biological labels often admit ontology-consistent descriptions at different levels of specificity. For example, “B cell” and “naive B cell” occupy an ancestor–subtype relation, whereas “B cell” and “neuron” belong to different branches. Exact string matching assigns the same zero reward to both cases and discards this biological structure. We therefore use a training-time Answer Judge that maps the prediction *ŷ* and target *y*^∗^ to a graded Cell Ontology-aware score [48]. This reward-side judge is distinct from the offline judge used for reported evaluation metrics (§4.4).

The Answer Judge assigns one of seven ordered semantic categories, ranging from exact or more-specific matches through valid ancestors to wrong-branch, wrong-lineage, and unrelated predictions. Its continuous score *s*_raw_ ∈ [−1, 1] is decomposed into 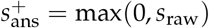 and 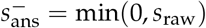. A separate Reasoning Judge is evaluated only after the answer enters the positive branch.

Figure 3 depicts the task-reward pipeline and the separation between answer semantics, reasoning quality, and structural safeguards.

**Figure 3.**
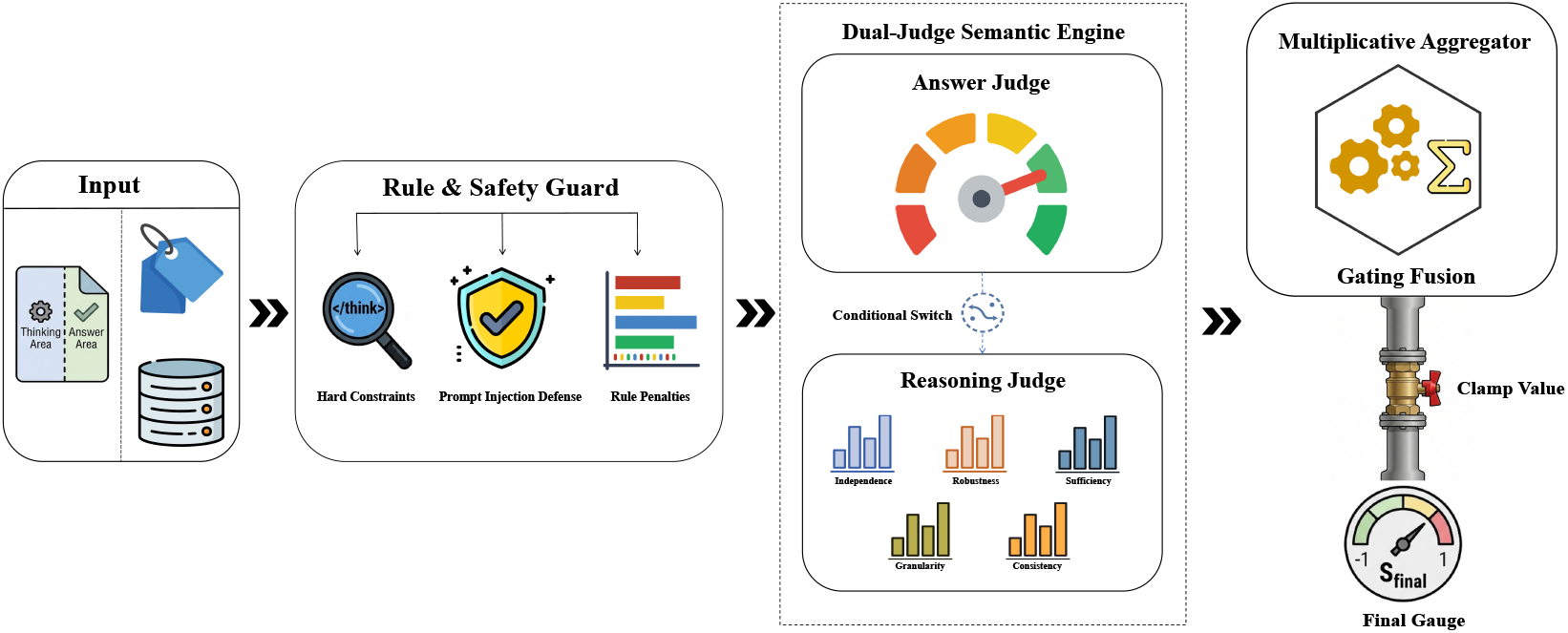
Ontology-grounded task-reward pipeline. Model outputs first pass through structural guards and then enter an Answer Judge that scores ontology-aware semantic correctness. Responses in the positive-answer branch are additionally assessed by a Reasoning Judge over evidence use, robustness, causality, granularity, and consistency. The resulting task reward is subsequently combined with the target-isolated factuality adjustment in Eq. 9.

##### Answer-gated reasoning credit

We factor answer correctness as a gate over reasoning credit:

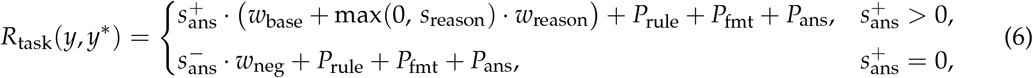

where *w*_base_, *w*_reason_, and *w*_neg_ control the answer and reasoning contributions, and *P*_rule_, *P*_fmt_, and *P*_ans_ are non-positive structural penalties. The gate has three consequences. An incorrect answer receives no reasoning credit; a correct answer benefits only from a positive reasoning score; and negative answer evidence retains its sign even when formatting or structural checks fail. For population tasks, per-cell semantic scores are aggregated before entering the same gate, with population-consistency terms defined in Appendix E.

##### Conditional reasoning evaluation

The Reasoning Judge is invoked only when the answer has positive semantic support and the structural preconditions pass. This confines reasoning credit to trajectories that reach an ontology-consistent conclusion. Its score is

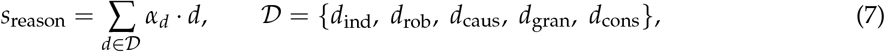

where the dimensions measure independence from the prompt, counterfactual robustness, causal sufficiency, granularity discipline, and internal consistency. Independence and robustness receive the greatest weight because they most directly test whether a rationale derives its conclusion from gene evidence and remains coherent under changes in biological context. Label Semantic Match is retained as a monitoring dimension but excluded from *s*_reason_ because the Answer Judge already scores that relation.

#### 3.4.2 Target-Isolated Atomic-Claim Verification

The task reward establishes whether a response reaches an ontology-consistent answer and uses a well-formed reasoning structure. Fact-Aware RLVR adds a complementary requirement: each biological assertion in that structure must remain traceable to evidence available for the analyzed cell or population.

##### Claims as auditable units

An extractor proposes minimal semantic claims and their sentence anchors, after which a deterministic compiler fixes the textual span, cell scope, gene identity, and dependencies of each claim. Admission requires an unambiguous source span, an eligible claim type and scope, content-level deduplication, and explicit dependency tracking. This converts a fluent rationale into a set of localized units that can be evaluated independently without allowing one supported statement to lend credit to another.

##### Three verification axes

Each admitted claim is evaluated along three axes: *truth*, whether expression or curated knowledge supports it; *grounding*, whether the cited evidence was visible to the model; and *overclaim*, whether an RNA-level observation has been promoted to a protein-, pathway-, functional-, or causal conclusion. Consider a claim that a marker gene is elevated and therefore establishes a cellular function. Its expression component may be supported on the truth axis, the grounding axis separately records whether the gene was available in the model-visible profile, and the overclaim axis determines whether the functional conclusion exceeds what RNA evidence licenses. Keeping the axes distinct preserves the difference between contradiction, inaccessible evidence, and an inference whose modality or causal scope is too strong.

The verifier combines the cell’s RVQ-decoded profile with frozen marker, regulon, pathway, and ontology resources. Each claim resolves to Supported, Contradicted, or Uncertain. Missing catalog entries, rank-truncated genes, and evidence withheld by mixed-lineage, dissociation, or cell-cycle guards resolve to Uncertain; they do not create contradictions.

##### Target-isolated evidence

The verification context contains the model-visible evidence and the source truth needed to check the claim, while excluding the label scored by the Answer Judge. Immutable evidence snapshots keep claim verification separate from target recovery, and an unresolved lookup returns Uncertain. This isolation lets the task reward assess the answer while the factuality lane assesses the evidence claims supporting it.

##### Bounded contradiction adjustment

Only unique, policy-eligible claims whose truth axis is Con-tradicted enter the factuality adjustment. Writing *n*_C_ for their count, distinct from the tokenizer’s *N*_*c*_ codebooks, we define

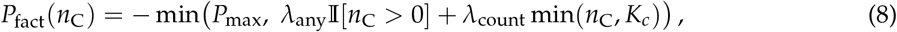

so that *P*_fact_ ∈ [− *P*_max_, 0]. The fixed term charges the presence of an eligible contradiction, the count term distinguishes repeated contradictions up to *K*_*c*_, and *P*_max_ bounds their total influence. Supported and uncertain claims contribute zero, making the adjustment a conservative penalty rather than an additional source of positive reward. Appendix E reports the active weights and verification specification.

The final reward clips the sum of task quality and factual consistency:

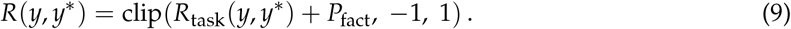

##### Group-level atomicity

GSPO compares sibling responses through the group-normalized advantages of Eq. 14, so their factuality adjustments must be computed under the same verification conditions. RVQ-Alpha therefore commits *P*_fact_ atomically across each sampled group: if any sibling is ineligible or any verification component fails, the group receives no factuality adjustment. Claim coverage and the uncertain fraction are retained as diagnostics and do not alter the reward.

### 3.5 MOPD: Multi-Teacher On-Policy Consolidation

Task-specific RLVR allows each policy to adapt within a homogeneous reward regime, producing specialists that exceed direct mixed-task training on the observed training families (Figure 4). A unified deployment target then requires one policy that can recover these complementary capabilities without checkpoint selection at inference time. Following MOPD in MiMo-V2-Flash [22] and related specialist-to-generalist distillation in Baichuan-M3 [23], RVQ-Alpha separates specialist formation from capability consolidation.

**Figure 4.**
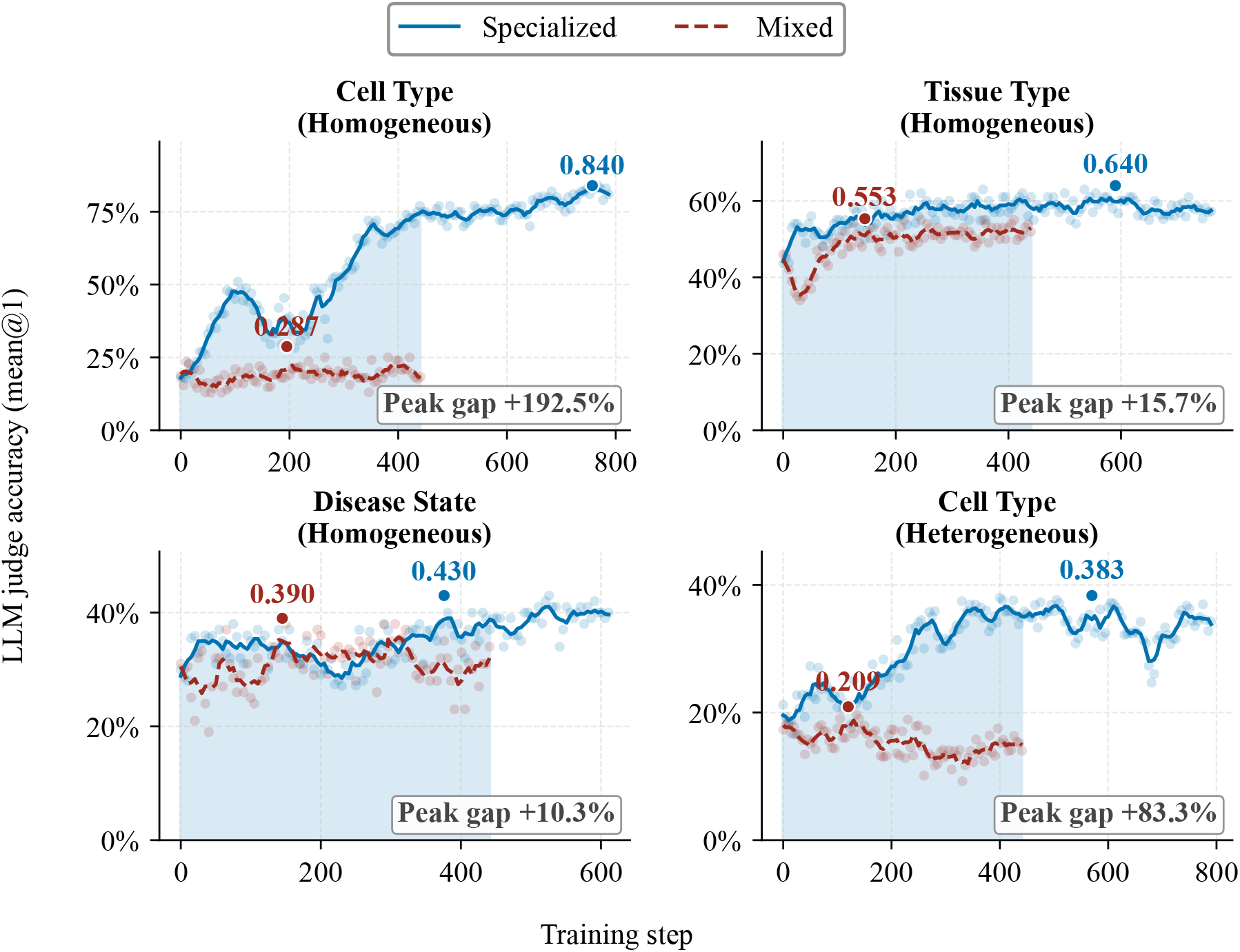
Specialized versus mixed-task RLVR training. Task-specialized policies attain higher judge accuracy than direct mixed-task training across the four observed training families. The gap motivates a separate consolidation phase in which routed specialist feedback is transferred into one student policy.

#### Specialist formation

Starting from the shared Evidence-First SFT checkpoint *π*_0_, we train *K* task-specialized policies, or specialists, 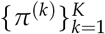 independently with GSPO:

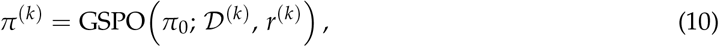

where *D*^(*k*)^ and *r*^(*k*)^ are the data and reward associated with route *k*. Once frozen for consolidation, each specialist serves as the teacher for its route. The pool covers single-cell cell-type identification and general-knowledge retention; homogeneous and heterogeneous population cell type; and homogeneous tissue type and disease state. A mixed-task GSPO checkpoint initialized from the same *π*_0_ provides the single-checkpoint baseline but does not serve as a teacher. This separation lets each teacher acquire its task competence before the student reconciles their output distributions.

#### On-policy task-routed consolidation

The student learns on trajectories generated by its own evolving policy. For each sample *x*, a deterministic router *τ*(*x*) selects the frozen teacher corresponding to its route. At sampled token *y*_*t*_, the clipped student–teacher log-ratio defines the distillation advantage

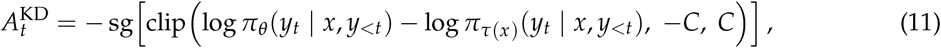

where sg[·] stops gradients through the advantage. The student is updated with a PPO-clipped surrogate over the same on-policy samples:

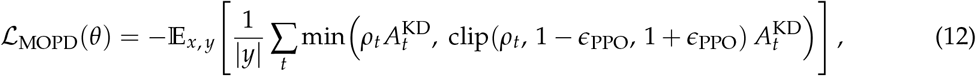

with *ρ*_*t*_ = *π*_*θ*_ (*y*_*t*_ | *x, y*_<*t*_)/*π*_old_(*y*_*t*_ | *x, y*_<*t*_). Because the teacher evaluates tokens sampled from the student’s current policy, the distillation signal follows the states the student actually visits instead of relying on a fixed archive of teacher demonstrations. The adopted route uses the distillation advantage alone in the update; task and judge scores monitor consolidation and retained-task performance without entering the distillation gradient.

#### Progressive consolidation across task routes

Capability consolidation proceeds through four additions:

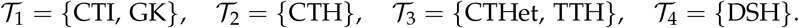

Here *T*_*r*_ denotes the routes newly admitted in round *r*, not the exclusive contents of its training batches. Each round initializes from the preceding student, routes new tasks to their specialists, and replays retained tasks through the previous consolidated teacher:

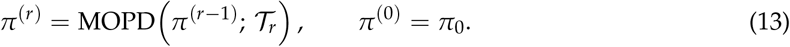

This progressive schedule turns the preceding student into the retention path for capabilities already consolidated, while newly admitted specialists supply task-specific guidance. The final unified checkpoint *π*^(*R*)^ is evaluated unchanged across the CAPSTONE tasks; the concrete teacher identities, checkpoint-selection schedule, and optimization settings are reported with the experimental protocol and reproducibility details.

## 4 Experimental Setup

The empirical study asks four questions: whether a compact cellular alphabet preserves biological structure; how capability changes across CAPSTONE shifts; whether specialist policies can be consolidated; and which components and training behaviors explain the resulting profile. We define the checkpoint lineage, biological exposure, and grading contract before reporting the corresponding evidence.

### 4.1 Controlled Backbone and Post-Training Regimes

#### Shared backbone and checkpoint lineage

All RVQ-Alpha variants use **Qwen3-4B-Thinking-2507** [49], a 4.0 B-parameter model with *V*_text_ = 151,941. We extend its vocabulary with *N*_*c*_ · *C* = 8 × 32 = 256 RVQ single-cell symbols, five structural delimiters, and the top-500 gene-name tokens (§3.2). The Thinking variant aligns the backbone’s native response regime with an RLVR objective that evaluates both biological conclusions and their reasoning traces (§3.4.1). Holding this backbone fixed makes post-training strategy, rather than architecture or parameter count, the principal variable within the RVQ-Alpha comparison.

The checkpoint lineage proceeds from continued pretraining (CPT) to Evidence-First supervised fine-tuning (SFT), after which the shared SFT checkpoint branches into direct mixed-task RLVR and independently optimized specialists. Multi-Teacher On-Policy Distillation (MOPD) then uses the frozen specialists as routed teachers and consolidates their capabilities into one student (§3.5). Thus, the four rows compared within the *Ours* block have distinct experimental roles: SFT is the common post-training origin, Mixed-Task is the direct unified-policy baseline, Specialist is a task-routed multi-checkpoint reference, and MOPD is the consolidated single checkpoint.

Table 2 summarizes the adaptation stages before MOPD consolidation. CPT and SFT each use four NVIDIA H20 96 GB GPUs; every RLVR run uses eight training GPUs, with eight additional GPUs serving the reward judge.

#### Continued pretraining

CPT establishes the initial association between the expanded vocabulary and biological text using a six-source mixture of ≈ 225,000 records combined in natural proportions. Eight donors (A013, D094–D096, D099, D101–D103) are reserved from this stage for donor-level evaluation. Packed sequences contain 16,384 tokens and are sharded with fully sharded data parallelism under a (data-parallel, context-parallel, tensor-parallel) degree of (2, 2, 1). AdamW optimization uses a peak learning rate of 5 × 10^−5^, 10% cosine warmup, and weight decay 0.1 under BF16 computation with FP32 gradient reduction and optimizer states. Across 13,000 updates at 32,768 tokens per step, the model observes approximately 4.26×10^8^ tokens, or one to two effective passes over the mixture.

#### Supervised fine-tuning

Evidence-First SFT turns the broad cellular–language associations acquired during CPT into task-conditioned biological responses. It initializes from the merged CPT checkpoint and uses ≈121,000 chat-template conversations with chain-of-thought reasoning from five sources while preserving the donor-held-out split. Supervision is restricted to assistant spans under the Qwen3 chat template. To retain the alignment established during CPT, SFT uses a light adaptation regime: the peak learning rate decreases to 1 × 10^−6^ and the warmup ratio increases from 0.1 to 0.2. Parallelism, precision, sequence length, and effective batch match CPT, and training covers two epochs at 32,768 tokens per step.

#### RLVR

Specialist RLVR introduces answer- and evidence-conditioned feedback after SFT. The per-task runs described here cover homogeneous tissue, homogeneous disease state, and heterogeneous cell type. We use GSPO [47] with the GRPO group-relative advantage estimator [50]; its response-level importance ratio and objective are defined in Appendix D. Because that geometric-mean ratio concentrates with response length, the update uses tight asymmetric bounds of 3×10^−4^/4×10^−4^ and no separate KL or entropy regularization. The reward combines structural checks, ontology-aware answer assessment, reasoning evaluation, and the target-isolated factuality adjustment defined in §3.4.1–§3.4.2. Each optimizer step draws 128 trajectories from 16 prompts and eight rollouts at temperature 1.0; training runs for 2,000 steps with AdamW at 1×10^−6^.

#### Specialization and MOPD consolidation

Mixed-Task and all specialists begin from the same Evidence-First SFT checkpoint, isolating how homogeneous versus joint reward regimes shape post-training. Once specialist optimization is complete, the resulting policies are frozen and become route-specific teachers for the on-policy MOPD student. Consolidation follows the four progressive route additions of §3.5, using the preceding student as the retention teacher for capabilities already integrated. Counts used elsewhere describe different partitions of this system: the three runs detailed above are optimizer runs, the four curves in Figure 4 summarize observed training families, and the six routes in §3.5 define the teacher inventory. The final MOPD checkpoint is evaluated unchanged across all CAPSTONE headline tasks.

#### Unified Checkpoint Protocol

To make stage comparisons less sensitive to a single noisy validation snapshot, CPT, SFT, and RLVR use one checkpoint-selection rule. Checkpoints are saved every 1,000 updates for CPT and SFT and every five optimizer updates for RLVR, scored on a fixed held-out benchmark, and the five highest-scoring checkpoints are averaged uniformly in parameter space. CPT and SFT are ranked by held-out single-cell LLM-judge accuracy, whereas each specialist is ranked by its task-specific validation accuracy. The merged checkpoint becomes the initialization for the next phase and the artifact used in the corresponding comparison.

All reported training regimes use seed 42. Checkpoint ranking relies on one stochastic decode per example (*T*=0.6, top-*p*=0.95, top-*k*=20), so the top-five merge is intended to reduce sensitivity to sample-level decoding noise rather than to estimate run-to-run variance. Full hyperparameter grids, prompt templates, mixed-precision settings, and checkpoint-merging details are provided in the supplementary protocol.

### 4.2 Datasets

#### Training corpus and biological breadth

We derive CPT and SFT examples from donor-level human scRNA-seq profiles in ScBank. The working corpus is a **55-file indexed subset** containing **31**,**481**,**661 cells** from approximately 45 source-study families. After metadata harmonization, it spans **101 standardized tissues, 103 disease states**, and **859 standardized cell-type labels** (Table 3). These figures describe the biological breadth of the harmonized corpus rather than the effective label count of any one downstream task. Fifty-three of the 55 dataset identifiers are linked to upstream ScBank source records; the remaining two are retained with their provenance in the released corpus index.

#### Harmonization and expression processing

Tissue, disease, and cell-type annotations are canonicalized through field-alias normalization and dictionary-based value mapping before task construction. Because metadata coverage varies by field—donor identifier and tissue are present in all 55 files, whereas disease, cell type, sex, and age have progressively lower coverage—we characterize biological exposure through the harmonized label space rather than assume a uniformly complete schema. Standard per-cell QC fields such as detected-gene count, total count, and mitochondrial fraction are sparse in the processed files; the corpus description is therefore anchored to metadata-supported harmonization. Expression binning and assignment to RVQ symbols are specified independently in §3.2.

#### Derived training pools

SFT and RLVR consume task-specific conversation records derived from the indexed corpus. Single-cell annotation uses a filtered pool of 98,873 cells for the main experiments; larger unfiltered pools of 286,452 and 2,867,679 cells are used only when explicitly identified. Counts reported for population tasks denote their construction pools, with task-level sampling applied when the corresponding training instances are formed.

### 4.3 Benchmarks

Downstream evaluation uses **CAPSTONE** (*Cell Annotation and Population-level tasks under Shift: a Testbed with Ontology-Normalized Evaluation*). Its four tasks ask complementary generalization questions at increasing biological scale: **CTA** assigns a cell type to a single cell; **TI** and **DSA** infer tissue and disease state from sample-level cell populations; and **PCA** reconstructs a donor-level cell-type composition. Each task pairs its biological unit with a shift axis, so the suite tests transfer across tissue, dataset, disease, or study context rather than merely repeating an in-distribution label lookup.

#### Two-level holdout

CAPSTONE separates source membership from task-level biological isolation. The indexed training inventory contains 55 source identifiers and the holdout layer contains 13 canonical dataset identifiers, with zero intersection after identifier normalization.

Within that source boundary, CTA holds out donors under context shifts; TI Core-OOD is study-disjoint; the eligible DSA Core-OOD endpoint follows leave-one-study-out evaluation; and PCA isolates donors under unseen-tissue or unseen-disease shifts. Evaluation cells are encoded by the frozen RVQ tokenizer without parameter updates. D102 is assigned exclusively to diagnostic analysis and contributes no headline record. Throughout the paper, *held-out evaluation* denotes these 13 CAPSTONE datasets under their task-specific isolation rules. The cell-type vocabulary is shared with training, so CTA measures transfer under biological shift rather than label-space novelty.

#### Task and split inventory

Table 4 enumerates each split’s protocol role, independent biological units, and reported evaluation records. CTA contains 3,840 cells across 264 Cell Ontology labels. TI, DSA, and PCA use nested population sizes *N* ∈{8, 16, 32, 64} where sufficient cells are available; consequently, several records may describe different population views of the same independent sample.

The headline comparison uses CTA cross-disease, TI Core-OOD, the 120-record DSA Natural Core-OOD scope, and an equal-weight macro over the two PCA axes. CTA cross-dataset and the DSA composition-matched and homotypic constructions remain diagnostic views rather than additional headline estimates.

#### Shortcut controls

Each headline axis includes a deliberately expression-free control. CTA uses a tissue-conditioned training-frequency prior; TI uses a metadata-only label prior estimated from training studies; and DSA uses a constant-prevalence predictor. These controls test whether the declared shift still permits the target to be recovered from contextual prevalence alone. Their outcomes are reported alongside the model results in §5.1.

### 4.4 Deterministic Evaluation Protocol

All headline scores in Table 5 come from deterministic, ontology-aware graders. Every system is evaluated on the same biological instances and gold targets with a shared split manifest, parser, and grader; model-based judging is reserved for diagnostic analyses.

#### Primary and complementary metrics

Macro-F1 is primary for CTA and TI, endpoint-macro AUPRC for DSA, and strict Jensen–Shannon divergence for PCA. Coarse-v1—a frozen, versioned, content-hashed label transform—is applied to both PCA gold compositions and predictions before divergence, placing donors annotated at different granularity on one common vocabulary. Exact Cell Ontology accuracy and pure *is-a* distance complement CTA; balanced accuracy and UBERON semantic similarity [51] complement TI; AUROC and balanced accuracy complement DSA; and detection AU-ROC together with strict dominant-type accuracy complement PCA. These separate exact agreement, ontology proximity, class-balanced discrimination, and distributional recovery.

#### Ontology pinning

Predictions are resolved against version-pinned releases of the Cell Ontology, UBERON, and MONDO. Release identifiers and graph hashes accompany each evaluation. If a pinned *is-a* graph is cyclic or does not contain a gold identifier, the run is rejected instead of silently falling back to a looser semantic match.

#### Failure handling

Malformed, unresolved, abstained, and missing predictions remain in the canonical denominator under a task-defined worst-case rule. CTA and TI count unresolved labels as incorrect; DSA retains invalid bag responses as non-successes during endpoint aggregation; and PCA assigns worst-case strict JSD and dominant-type accuracy while treating the corresponding composition as empty for detection AUROC. No failure category is used as an exclusion rule.

#### Diagnostic judge

The offline judge supplies only the per-cell-type and misclassification-flow analyses in App. A–App. B; none of its outputs enters Table 5. Following Zheng et al. [52], DeepSeek-chat assigns a task-aware semantic relation and a score in [0, 1]. Diagnostic accuracy uses *τ*=0.8 for single-cell CTA, *τ*=0.75 for cell-type and tissue population views, and *τ*=0.85 for disease-state populations. Non-leaf accuracy additionally credits valid subtypes of the gold label, while weighted F1 complements macro-F1 under class imbalance.

This diagnostic stack differs from the Qwen3-30B-A3B reward and monitoring stack in model family, endpoint, prompts, and aggregation. The separation reduces direct reuse of the training judge and its attendant self-preference risk [53]; it does not imply formal statistical independence. Reward-side sampling and aggregation are specified in App. E.

#### Split integrity and run provenance

At the representation level, the 13 canonical holdout identifiers are absent from RVQ pretraining; at the supervision level, task builders isolate donors, studies, or endpoints according to the rules above and group population records by sample-processing identifier. CAPSTONE does not independently certify upstream donor closure beyond these dataset- and task-level checks. Each evaluation records its decoding regime, serving provider, resolved model path, input manifest, runtime, and code commit. Content hashes bind the metric-bearing inputs and outputs, while benchmark-generation code, sampled evaluation sets, and reference configurations support re-evaluation under the same contract.

### 4.5 Baselines

Table 5 includes systems whose released native workflow produces an output that the CAPSTONE grader can score without an additional learned adapter or task-specific fine-tuning. Model scale is reported as an interpretive variable rather than used as an inclusion filter.

The general-LLM block covers Qwen3 (4B-Thinking, 30B-A3B-Thinking, and 35B-A3B), Kimi-K2, GPT-OSS (20B and 120B), Doubao-Seed-2.0-Pro, and DeepSeek-V4 (Flash and Pro). The Qwen3 models receive ranked gene-name serializations of the same biological instances and therefore serve as text-interface controls across three scales. The domain block covers Cell-o1 [18], CellTok [19], C2S-Scale-Gemma-2-2B [13], and CellWhisperer [35]. Unsupported task–system pairs remain “—” rather than being populated through an improvised bridge.

Systems whose native workflow emits no task-compatible label remain outside the table: scGPT [5] and Geneformer [6] produce cell embeddings and would require an external protocol bridge, and CellFM [25] requires task-specific fine-tuning. App. G gives the per-system rationale. Commercial-model training exposure cannot be audited, so those rows remain descriptive.

## 5 Results

The empirical comparison reveals three recurring patterns. Capability rankings change with the biological question and the metric used to score it; post-training produces a consistent progression within the controlled RVQ-Alpha family; and specialist consolidation recovers most, but not all, of the task-routed frontier. We first establish these cross-system profiles, then examine consolidation and the component-level probes that qualify their interpretation.

### 5.1 Task- and Metric-Dependent Capability Profiles

Table 5 scores every system on the four CAPSTONE headline views under one deterministic grading contract (§4.4). All systems see the same biological instances, gold targets, split manifests, parsers, and graders; no headline record in this table entered post-training or checkpoint selection.

The strongest model depends on what aspect of cellular reasoning is emphasized. The 4.0 B MOPD checkpoint reaches CTA macro-F1 0.328, TI macro-F1 0.674, and PCA detection AUROC 0.775, narrowly exceeding the best external values of 0.325, 0.668, and 0.769, respectively. The ordering reverses elsewhere: Cell-o1 attains DSA AUPRC/AUROC 0.615/0.810, compared with 0.604/0.796 for MOPD, while DeepSeek-V4-Pro reaches PCA strict JSD/dominant-type accuracy 0.045/0.976, compared with 0.251/0.972. CAPSTONE therefore exposes a task- and metric-dependent capability profile rather than a single global ranking. Cross-family contrasts remain descriptive because model scale, training exposure, and input interface differ among systems.

#### Text-input controls

Moving from the 4B Qwen text-input control to the 30B and 35B variants sharply improves CTA, TI, and PCA, but scale does not induce a uniform ordering across PCA metrics. The 30B model has lower strict JSD and higher dominant-type accuracy than the 35B model, whereas the 35B model has higher detection AUROC. Scale therefore changes the capability profile without producing a uniform ordering across population metrics.

#### Controlled comparison within RVQ-Alpha

Within the *Ours* block, all regimes share one backbone and SFT origin, making the effect of post-training directly visible. The primary metrics progress from SFT to Mixed-Task to MOPD: CTA macro-F1 rises 0.281 → 0.309 → 0.328; TI macro-F1, 0.569 → 0.634 → 0.674; DSA AUPRC, 0.491 → 0.553 → 0.604; and PCA strict JSD falls 0.342 → 0.287 → 0.251. The same ordering holds for all twelve reported metrics. SFT, Mixed-Task, and MOPD are each evaluated as one unchanged checkpoint, whereas the Specialist row selects a task-matched checkpoint and serves as the within-family specialization reference.

#### Population-level scope

TI, DSA, and PCA extend the interface from isolated cells to nested bags containing up to 64 cells. Population rewards aggregate per-cell judgments while tolerating a bounded number of missing or underdeveloped cells (Appendix E), discouraging reliance on a single salient cell. These experiments concern sampled cell populations rather than atlas-scale inference.

#### Expression-free stress tests

Expression-free controls distinguish biological transfer from contextual prevalence. The CTA tissue-frequency prior falls from macro-F1 0.070 in distribution to 0.017 across tissue, 0.003 across dataset, and 0.000 across disease. On TI Core-OOD, the metadata-only predictor scores macro-F1/balanced accuracy/UBERON similarity 0.000/0.000/0.753 despite remaining strong within study, supporting the use of the study-disjoint view as the headline. The DSA constant-prevalence floor yields AUPRC/AUROC/balanced accuracy 0.333/0.500/0.500. Together, these controls show that the reported shifts substantially weaken simple contextual priors.

### 5.2 MOPD Nearly Closes the Specialist Gap

Figure 4 establishes the specialization frontier that motivates consolidation: direct mixed-task RLVR remains below the corresponding specialized trajectories across all four observed training families. Table 5 then asks how much of that frontier can be recovered without task-specific checkpoint routing.

MOPD improves over Mixed-Task on all 12 metrics. Relative to Mixed-Task it raises CTA and TI macro-F1 by 6.2% and 6.3%, raises DSA AUPRC by 9.2%, and lowers PCA strict JSD by 12.5%; in absolute terms the four primary metrics move by +0.019, +0.040, +0.051, and −0.036.

MOPD nevertheless remains slightly below the task-routed Specialist row on all 12 metrics. The four primary-metric gaps are 0.001 for CTA macro-F1, 0.002 for TI macro-F1, 0.003 for DSA AUPRC, and 0.005 for PCA strict JSD. Each lies within the predeclared 0.02 retention margin, so consolidation nearly closes, but does not erase, the observed specialist gap. This is a margin-based comparison from single-seed runs, not a statistical equivalence claim.

### 5.3 Grounding, Residual Refinement, and Input Dependence

Table 6 gathers three complementary probes: how grounding components alter legacy-holdout behavior, whether the residual cascade refines its estimate stage by stage, and whether recovered gene evidence depends on the paired cellular code.

**Table 6:** Ablations and controlled probes. (a) Single-seed component-removal deltas measured on the legacy holdout rather than on CAPSTONE; † denotes out-of-distribution accuracy, and *Severe* counts predictions lying more than five *is-a* edges from the ground-truth label. (b) Prefix decoding from a frozen tokenizer using only the first *K* codebooks; *Residual* is the fraction of latent residual energy remaining and *Pearson* is the per-gene correlation of the reconstruction. (c) True-minus-swap RVQ–gene pairing on top-50 gene recovery, with bootstrap intervals. Panels (b) and (c) are macros over twelve datasets. *Notes (qualitative failure modes)*. (i) **Without the specificity-penalty gate**, the model collapses to enumerating many gene markers without committing to a cell-type call. (ii) **Without the prompt-imitation detector**, 12% of generations contain near-verbatim system-prompt fragments; enabling the detector reduces this to <1%. Both behaviours are suppressed by the dual hard gates A′ and A′′ (Appendix E).

| (a) Core Component Removal |  |  |  |
| --- | --- | --- | --- |
| Removal | $\Delta\text{Acc. (pp)}$ | $\Delta\text{Halluc. (pp)}$ | $\Delta\text{Severe (\%)}$ |
| Evidence-First | -6.3 | +13.1 | — |
| Fact Verification | -3.2<sup>†</sup> | — | +12 |
| (b) RVQ Prefix Probe |  |  |  |
| $K$ | Residual ↓ | Pearson ↑ | |
| 0 | 1.000 | 0.442 |  |
| 1 | 0.211 | 0.575 |  |
| 2 | 0.165 | 0.603 |  |
| 4 | 0.130 | 0.623 |  |
| 8 | 0.095 | 0.636 |  |
| (c) RVQ-Gene Pairing (True – Swap) |  |  |  |
| Swap | $\Delta\text{F1@50} \uparrow$ | 95% CI | |
| Same type | +0.006 | $[-0.001, 0.014]$ | |
| Random | +0.043 | $[0.022, 0.066]$ | |

#### Component removal

Panel (a) reports single-seed component removal on the legacy holdout, rather than a repeated CAPSTONE estimate. Removing Evidence-First supervision—allowing a direct label without gene evidence—lowers accuracy by 6.3 pp and raises hallucination by 13.1 pp. Removing Fact Verification while retaining the other reward terms lowers OOD accuracy by 3.2 pp and increases severe ontology errors by 12%. Within this diagnostic scope, the two components leave distinct error signatures: Evidence-First supervision affects whether an assertion is grounded, whereas verification affects the biological distance of the remaining errors.

#### Residual prefix refinement

Panel (b) decodes from a frozen tokenizer using only the first *K* codebooks. The first codebook removes 78.9% of the initial residual energy and raises per-gene Pearson correlation from 0.442 to 0.575. Subsequent prefixes reduce the residual monotonically to 0.095 and raise correlation to 0.636, revealing a coarse initial estimate followed by progressively smaller refinements. This prefix probe is the quantitative evidence for residual refinement; Figure 5 provides a descriptive view of the accompanying code transitions.

**Figure 5.**
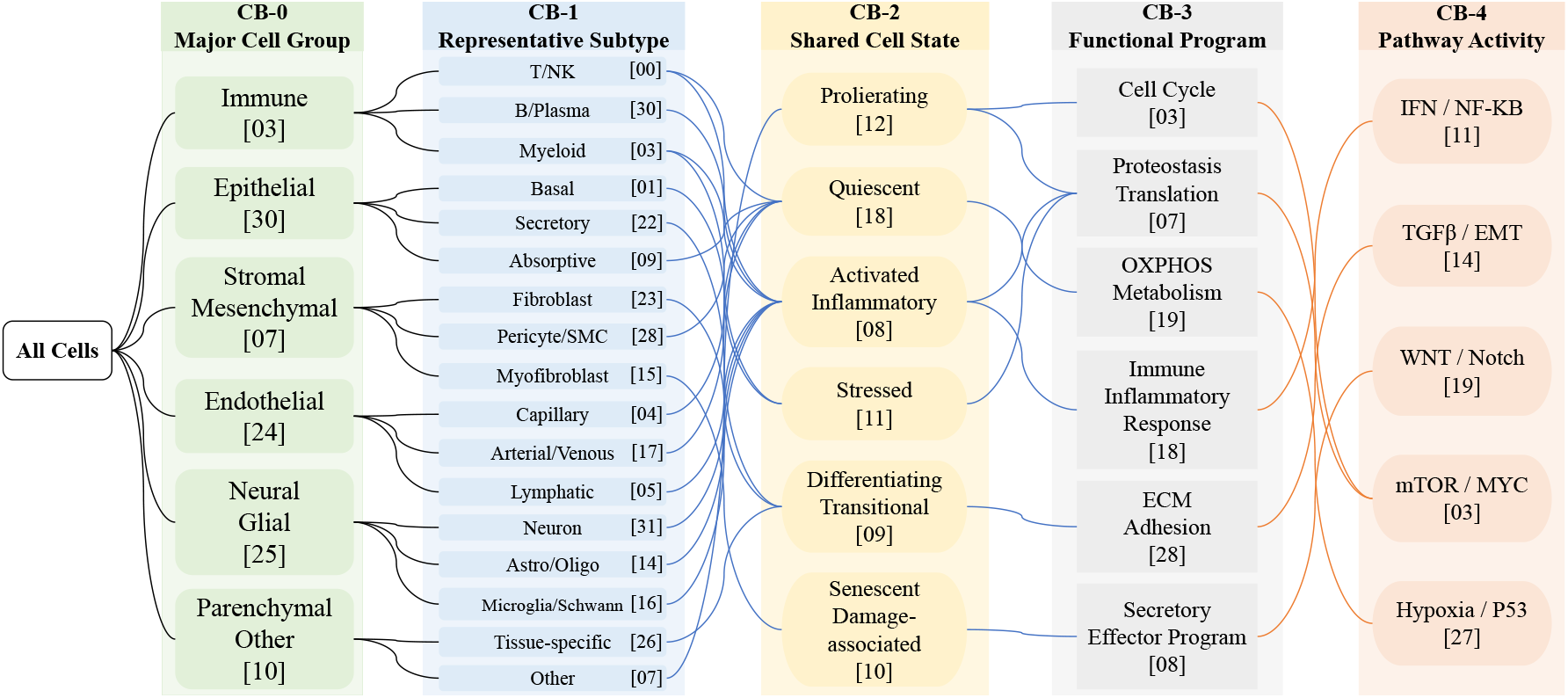
Empirical RVQ transition atlas across cell lineages. Directed transitions observed among 100K class-balanced cells encoded with (*N*_*c*_=8, *C*=32). The five displayed levels summarize associations with major cell groups, representative subtypes, shared states, functional programs, and pathway activity. Black edges denote the earlier branching view; blue and orange edges show the increasingly many-to-many transitions at later levels. The atlas is descriptive and does not establish intrinsic semantics or a biological ordering of the codebooks.

#### RVQ–gene pairing

Panel (c) swaps a cell’s RVQ codes for those of another cell and measures the true-minus-swap change in top-50 gene-recovery F1. Random pairing yields +0.043 with a 95% interval of [0.022, 0.066], supporting broad dependence of recovered genes on the input codes. The same-cell-type estimate is +0.006 with an interval that crosses zero, leaving cell-specific recovery within a shared type unresolved.

#### Reward-hacking defenses

The specificity gate suppresses the qualitative marker-stuffing pattern in which a response enumerates many markers without committing to a cell type. The prompt-imitation detector reduces near-verbatim system-prompt fragments from 12% to <1% of generations. Both defenses operate through the dual hard gates A^′^ and A^′′^ described in Appendix E.

## 6 Analysis

### 6.1 Compression–Structure Trade-off of the Cellular Alphabet

Table 7 compares six cellular alphabets parameterized by codebook count and size, (*N*_*c*_, *C*). The adopted (8, 32) configuration has the strongest observed joint profile: it leads seven of nine metrics, including reconstruction error (MSE 0.0738), distributional discrepancy (MMD 0.0298), per-cell correlation (0.704), cosine similarity (0.708), and all three clustering measures (ARI 0.247, AMI 0.557, NMI 0.589). It does not dominate every criterion. Usage is second at 52.31%, behind (32, 32) at 55.18%, and per-gene correlation is third at 0.116.

**Table 7:** RVQ tokenizer configuration ablation. Six (*N*_*c*_, *C*) variants evaluated on held-out cells, where *N*_*c*_ is the codebook count and *C* the codebook size. *Usage* is the fraction of active entries; PCC_*c*_/PCC_*g*_ are per-cell/per-gene Pearson correlations; ACS is average cosine similarity; MMD is maximum mean discrepancy; and ARI/AMI/NMI measure clustering against held-out cell-type labels. Bold marks the best value per column; ⋆ marks the adopted configuration, which leads seven of nine metrics among the six tested variants.

| $(N_c, C)$ | Usage % | MSE ↓ | $PCC_c$ ↑ | $PCC_g$ ↑ | ACS ↑ | MMD ↓ | ARI ↑ | AMI ↑ | NMI ↑ |
| --- | --- | --- | --- | --- | --- | --- | --- | --- | --- |
| $(8, 32)^*$ | 52.31 | **0.0738** | **0.704** | 0.116 | **0.708** | **0.0298** | **0.247** | **0.557** | **0.589** |
| $(8, 256)$ | 24.02 | 0.0775 | 0.690 | 0.113 | 0.692 | 0.0332 | 0.245 | 0.552 | 0.584 |
| $(16, 64)$ | 43.46 | 0.0771 | 0.694 | 0.115 | 0.698 | 0.0327 | 0.220 | 0.543 | 0.577 |
| $(16, 512)$ | 9.81 | 0.0796 | 0.697 | 0.117 | 0.702 | 0.0368 | 0.217 | 0.521 | 0.556 |
| $(32, 32)$ | **55.18** | 0.0759 | 0.695 | **0.118** | 0.698 | 0.0309 | 0.214 | 0.541 | 0.575 |
| $(128, 8)$ | 50.39 | 0.0801 | 0.652 | 0.105 | 0.661 | 0.0388 | 0.220 | 0.535 | 0.569 |

The alternatives reveal two opposing inefficiencies. Expanding the alphabet to (16, 512) activates only 9.81% of its 8,192 entries, leaving most nominal capacity unused. Extending the residual sequence to (128, 8) instead lowers per-cell correlation to 0.652, lowers cosine similarity to 0.661, and raises MMD to 0.0388 without a compensating clustering gain. Against these extremes, (8, 32) adds only 256 LLM vocabulary entries and represents each cell with eight symbols. Its selection therefore reflects the best observed compression–structure trade-off among the six tested configurations, rather than optimality on every individual metric.

### 6.2 Residual Refinement and Gene-Linked Evidence

#### Code-transition structure

Residual codebook position does not encode an intrinsic biological hierarchy: every stage is learned under the same reconstruction objective, and the transition atlas in Figure 5 is an empirical association map rather than a causal decomposition. Within that boundary, earlier observed transitions (CB-0–CB-2) align more often with broad identity groupings, representative subtypes, and shared cell states, whereas later displayed transitions (CB-3–CB-4) form many-to-many associations with functional programs and pathway markers. The prefix probe in §5.3, rather than the visualization alone, supplies the quantitative evidence that later stages refine an earlier estimate.

#### Input-dependent gene evidence

The pairing probe in Table 6 supplies a controlled test of whether recovered gene evidence depends on the paired cellular code. Random code swaps reduce gene recovery, whereas same-type swaps do not yield a resolved difference. The probe therefore supports input dependence without assigning intrinsic semantics to individual code positions; direct correspon-dence between those positions, attention, and specific gene modules remains a question for targeted mechanistic analysis.

### 6.3 Reward Dynamics and Task-Dependent Alignment

The training traces preserve a visible separation between the positive reward path 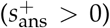 and the negative path under the piecewise objective of Eq. 6. The hard gates A^′^ and A^′′^ suppress the observed marker-stuffing, template-regurgitation, and multi-pattern over-specification behaviors; Table 6 quantifies prompt imitation and records the marker-stuffing failure mode.

Figure 6a follows response length and judge accuracy during cell-type RLVR. Length first rises during exploration, peaks near 9,500 tokens around step 250, and then contracts toward 7,000 tokens while accuracy continues to rise. The trace establishes a post-exploration contraction of the sampled responses; by itself, it does not identify the mechanism that produces shorter reasoning chains.

**Figure 6.**
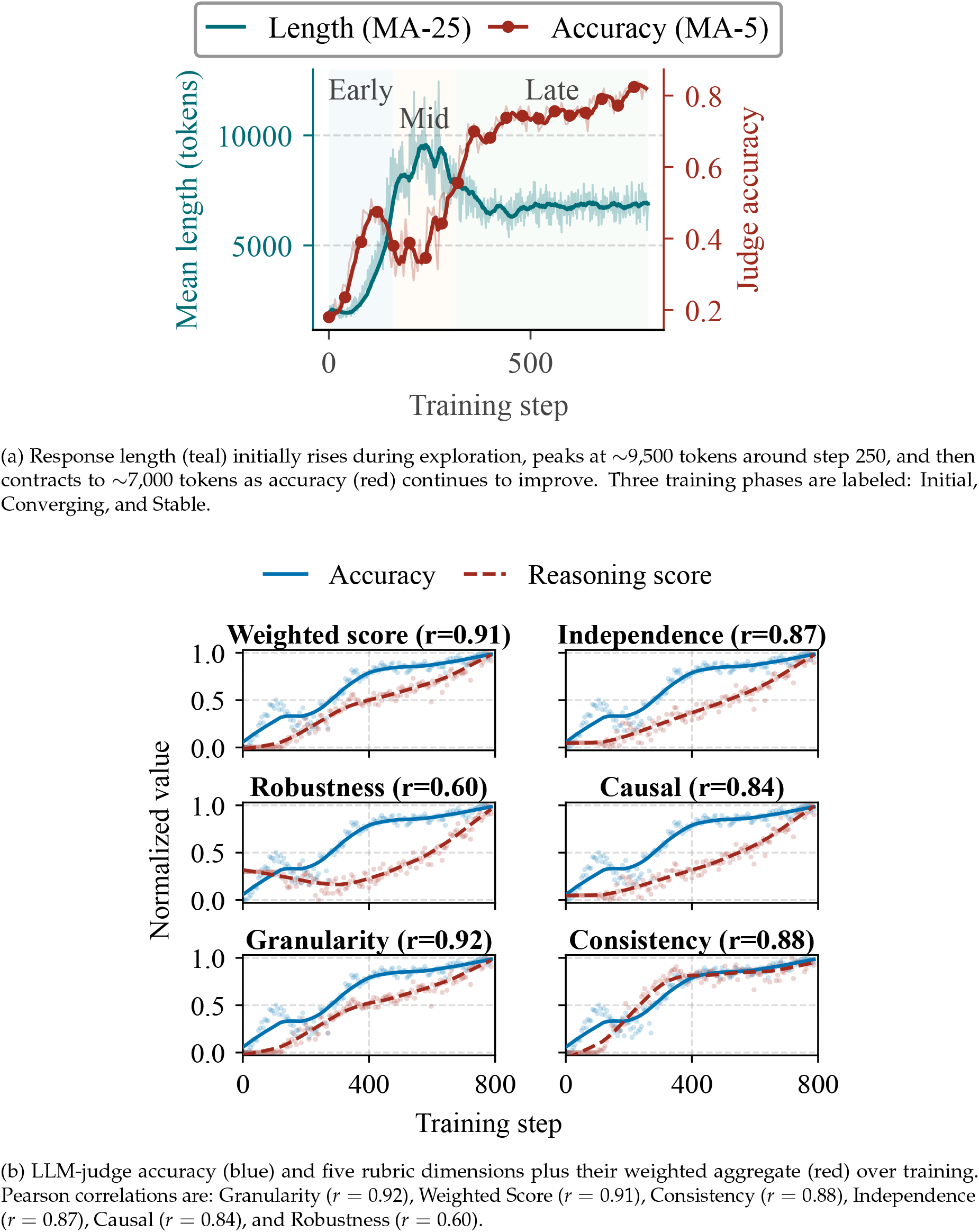
Reward dynamics for cell-type classification. (a) Response length and judge accuracy over training. (b) Co-movement between judge accuracy and rubric-based reasoning scores

Figure 6b compares the same judge accuracy with five rubric dimensions and their weighted aggregate. For cell-type training, these monitoring scores co-vary positively with accuracy (*r* = 0.60–0.92). Because both quantities are observed along one optimization trajectory and arise from related evaluation machinery, the correlations characterize co-movement rather than establish that the reward causally improves each reasoning property.

The corresponding tissue and disease traces (Supplementary Figures 11, 12, 13, and 14) expose an important task boundary. Correlations weaken from cell type (*r* ∈ [0.60, 0.92]) to tissue type (*r* ∈ [0.10, 0.74]) and disease state (*r* ∈ [−0.13, 0.70]); for disease state, Robustness is negatively correlated with judge accuracy. Reward alignment is therefore task-dependent: strong co-movement on cell identity does not imply that every reasoning dimension improves for phenotypically subtler endpoints.

## 7 Limitations

RVQ-Alpha presents several limitations that provide directions for future work:

### RVQ information loss

Compressing ~ 20,000 gene dimensions into 8 discrete tokens inherently discards fine-grained expression information. Although this representation is sufficient for cell type classification, the loss of information may degrade performance on tasks that require precise quantitative expression values (e.g., differential expression fold-change prediction).

### Evaluation scope

We report results on the 13 canonical CAPSTONE test datasets under donor-held-out, study-disjoint, and leave-one-study-out isolation (§4.3). Two residual limits remain. The DSA out-of-distribution headline rests on a single held-out endpoint, so its interval is correspondingly wide. And generalization to other dataset compositions, non-human organisms, and rare pathological states remains to be established.

### Computational cost

Each RLVR step issues multiple LLM-as-judge queries per generated response (three samples per response in our setup, adding approximately one second of latency per query). LLM judging and claim extraction together account for about 90% of RLVR training wall-clock time, which makes the reward loop the principal practical bottleneck. Distilling the judge into a lightweight classifier is the immediate engineering target.

### Gene name dependency

During training, Feature-Injection SFT relies on the top-*K* gene names. At inference time, without gene injection, the model must rely entirely on learned RVQ semantics; this dependence can degrade performance for very rare cell types that have limited training coverage.

### Population scale

Due to context length constraints, current population-level tasks operate on bags of up to 64 cells. Scaling to atlas-level populations (thousands of cells) requires efficient attention mechanisms or hierarchical summarization techniques.

### Baseline comparison scope

The cross-model comparison (Table 5) includes a system only where its released native workflow supplies a task-compatible output contract (§4.5). scGPT, Geneformer, and CellFM fall outside that criterion because they emit encoder-only outputs or require task adaptation; App. G discusses them without numerical evaluation.

### Baseline training-data overlap

Comparisons against closed or partially documented systems remain descriptive because their training exposure cannot be inspected. The training corpus of C2S-Scale-Gemma-2-2B is not fully documented [13], and the proprietary API models disclose neither corpus nor cutoff, so prior exposure to CAPSTONE source studies cannot be ruled out for those rows. CAPSTONE controls exposure for RVQ-Alpha itself through a two-level holdout with zero normalized dataset-ID overlap (§4.3).

### Single base model

We report all results using a single base model (Qwen3-4B-Thinking-2507, 4.0 B parameters; §4). We do not report variance estimates across random initializations; characterizing seed-level variance remains a direction for future work.

## 8 Conclusion

We presented RVQ-Alpha, an end-to-end framework that bridges single-cell transcriptomics and large language models through three key components: RVQ discrete tokenization, Evidence-First Feature-Injection SFT, and RLVR with fact verification. By converting continuous gene expression profiles into discrete tokens and embedding them directly into the LLM vocabulary, RVQ-Alpha places cellular evidence and text in one token stream, and a paired decoder maps those tokens back to the reference expression frame.

On CAPSTONE, MOPD improves over mixed-task training on all 12 metrics while retaining at least 99.5% of the task-routed specialists’ macro-F1 and AUPRC in a single deployable checkpoint (Table 5). The fact verification module grounds model outputs in actual expression data, reducing both hallucination and severe ontology errors (Table 6).

RVQ-Alpha establishes a foundation for AI Virtual Cells, providing a scalable and interpretable substrate for multi-modal biological intelligence. Future work will extend this framework to crossspecies generalization, atlas-scale populations, and additional modalities (spatial transcriptomics, chromatin accessibility). The next test for this interface is quantitative: expression-profile reconstruction and perturbation prediction.

## Supplementary Material

### Notation and scope

Two symbols are close enough to warrant disambiguation. Throughout, *N*_*c*_ denotes the number of RVQ codebooks in the tokenizer cascade (*N*_*c*_ = 8 in the deployed configuration), whereas *n*_C_ denotes the count of unique, policy-eligible contradicted claims entering the factuality adjustment of Eq. 8. The four CAPSTONE task abbreviations are CTA (cell-type annotation, single cells), TI (tissue identification, population bags), DSA (disease-state annotation, population bags), and PCA (population composition analysis, donor-level distributions); PCA in this sense is unrelated to principal component analysis.

### Roadmap

This appendix collects supplementary figures and tables referenced from the main text. App. A gives the retained cell-type misclassification-flow analysis (Figure 7); App. B the per-cell-type accuracy distributions (Figures 8, 9, 10); App. C the cross-task reward dynamics (Figures 11, 12, 13, 14); App. D the sequence-level policy objective and its distinction from token-level GRPO; App. E the reward-system design details (Tables 9, 10); App. F the RVQ tokenizer architecture, training, and vocabulary expansion, together with the codebook-utilization, prefix-*K*, token-efficiency, and cross-method diagnostics (Tables 11, 12, 13, 14, 15); App. G the per-system rationale for systems absent from the cross-model comparison; and App. H the model-interface positioning of CAPSTONE relative to existing single-cell evaluation resources.

**Figure 7.**
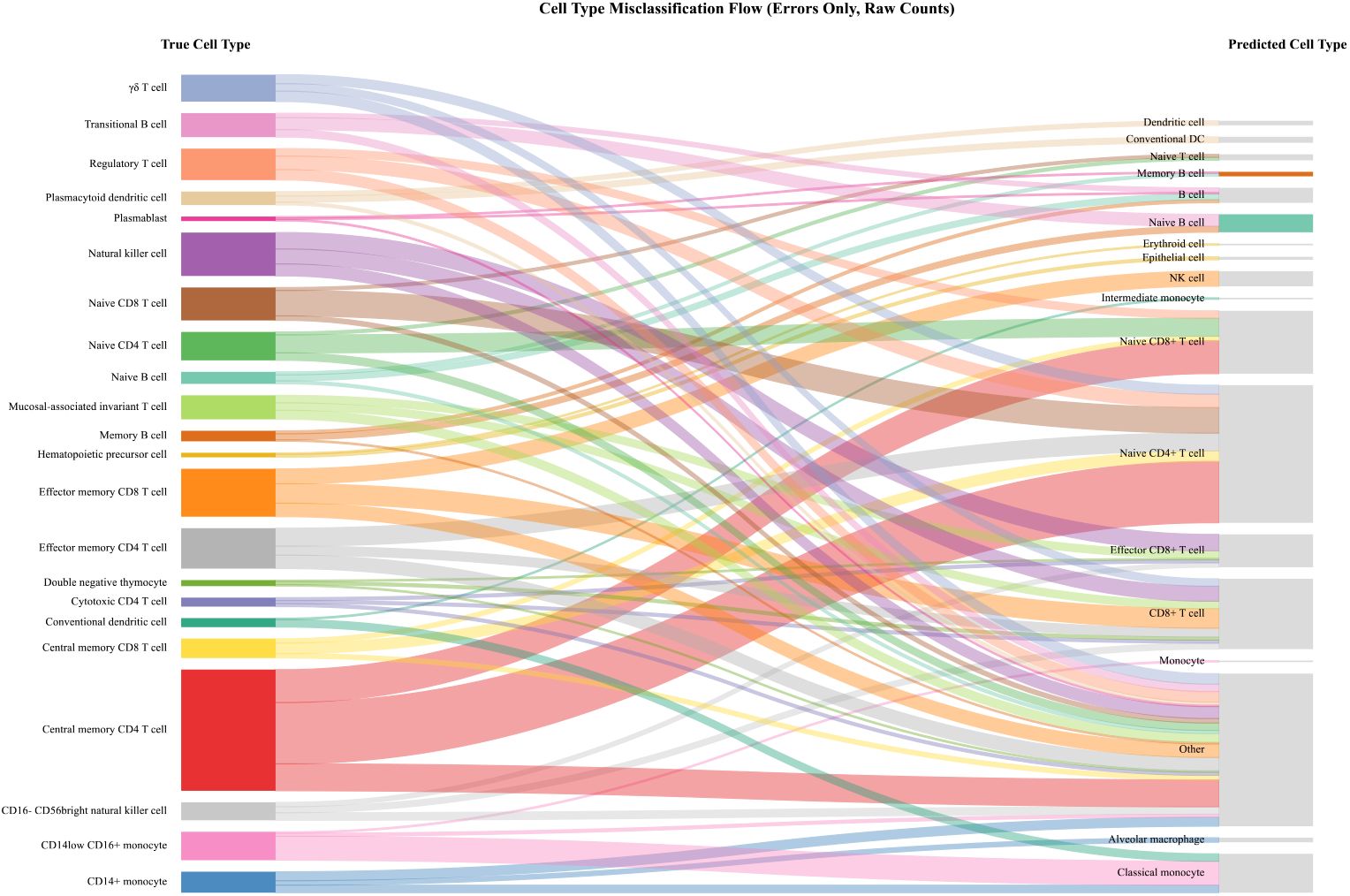
Misclassification flow by cell type. Two-column alluvial diagram mapping true cell types (left, 22 lineage-ordered nodes) to predicted cell types (right, canonicalized labels). Flow width scales as the square root of error count (offline evaluation judge threshold 0.8, the single-cell operating point of §4.4; top-2 destinations per true class). Predicted labels are normalized through 65 canonicalization rules that collapse them to 23 canonical forms. Central memory CD4 T cell generates the largest total error volume, with flows fanning out to Naive CD4+ T cell, Effector CD8+ T cell, and Monocyte. CD14+ monocyte produces the second-largest error mass, flowing primarily to Classical monocyte and Alveolar macrophage. Most T-cell subtypes flow to other T-cell predictions (sibling confusion), while cross-lineage errors (e.g., T cell → Epithelial cell) represent thin flow bands, confirming that ontologically close misclassifications dominate over cross-lineage errors.

**Figure 8.**
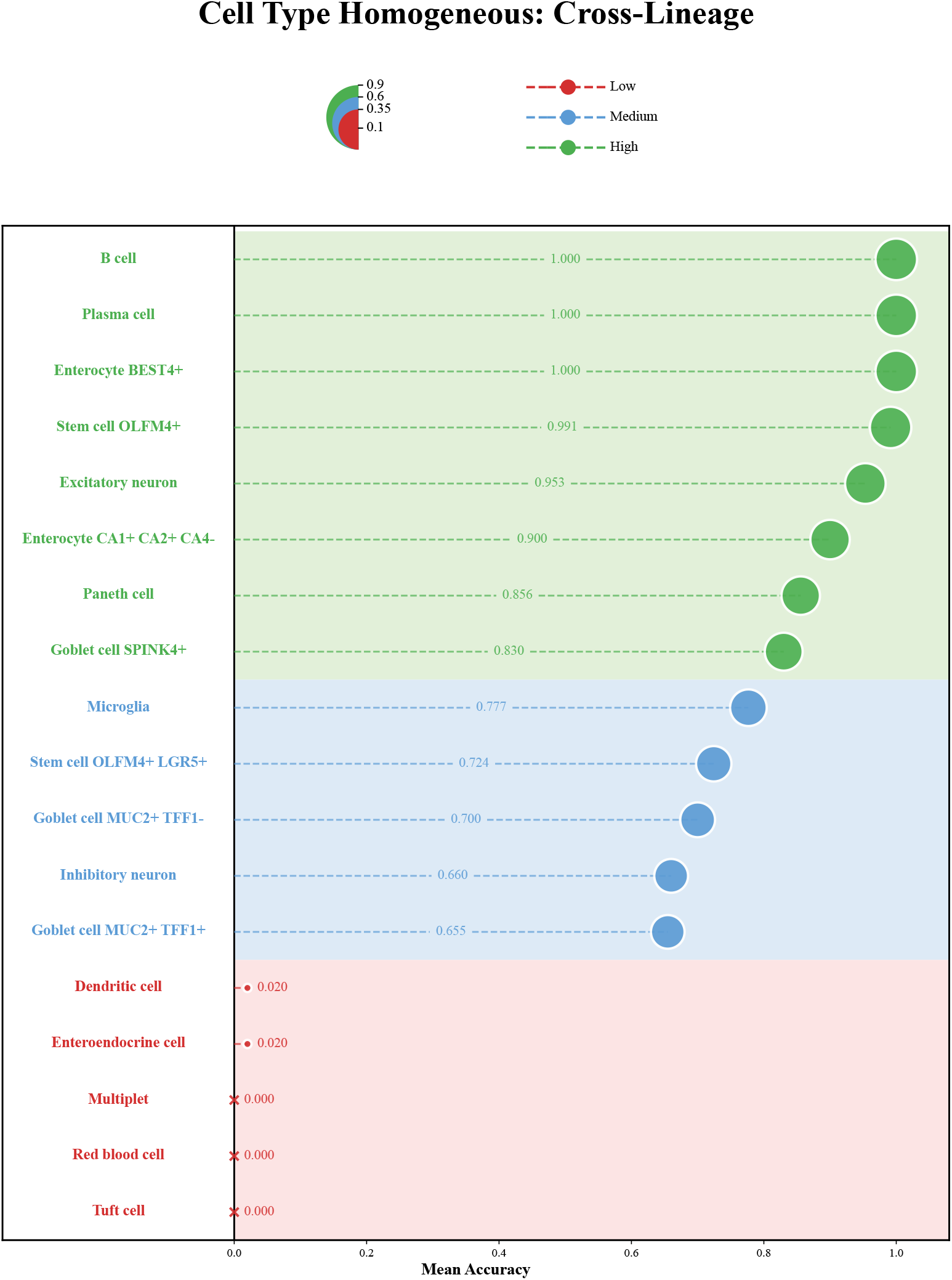
Per-cell-type accuracy distribution — cell type classification. Lollipop chart of per-cell-type accuracy on the homogeneous cell-type task (the converged GSPO cell-type policy; offline evaluation judge threshold 0.75; 9 lineage groups). **High tier** (≥0.8): B cell, Plasma cell, and Enterocyte BEST4+ all achieve 1.000, along with Stem cell OLFM4+ (0.991), Excitatory neuron (0.953), Enterocyte CA1+CA2+CA4 − (0.900), Paneth cell (0.856), and Goblet cell SPINK4+ (0.830). **Medium tier**: Microglia (0.777), Stem cell OLFM4+ LGR5+ (0.724), Goblet cell MUC2+ TFF1− (0.700), Inhibitory neuron (0.660), Goblet cell MUC2+ TFF1+ (0.655). **Low tier**: Dendritic cell (0.020), Enteroendocrine cell (0.020), and three zero-accuracy types (Multiplet, Red blood cell, Tuft cell). The long lower tail reflects rare and ontologically proximate cell types that the model struggles to distinguish.

**Figure 9.**
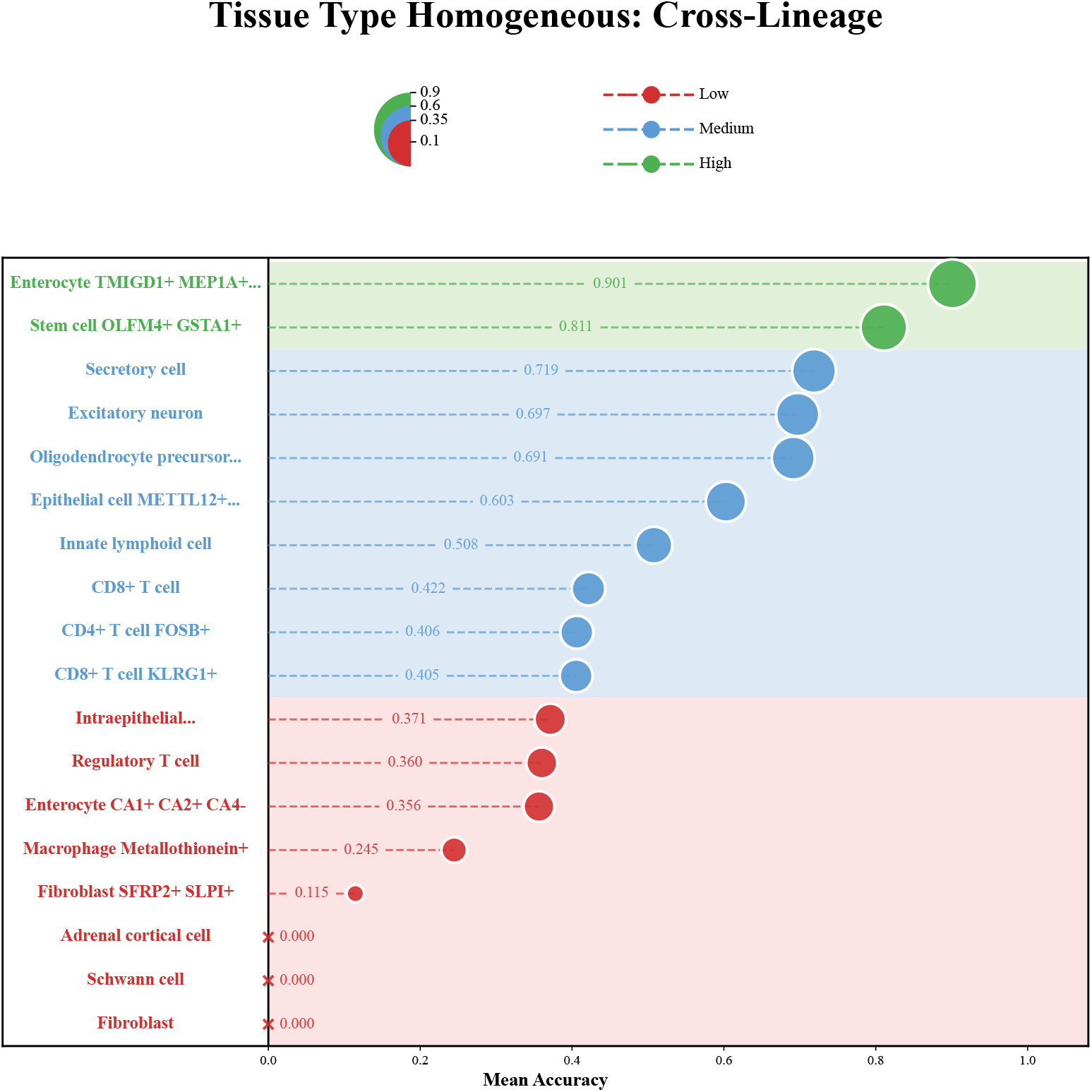
Per-cell-type accuracy distribution — tissue type classification. Lollipop chart of per-cell-type accuracy on the homogeneous tissue-type task (the converged GSPO tissue policy; offline evaluation judge threshold 0.75; 10 lineage groups including separate NK/ILC and Stromal lineages). **High tier**: only two cell types qualify—Enterocyte TMIGD1+ MEP1A+ (0.901) and Stem cell OLFM4+ GSTA1+ (0.811)—far fewer than the eight High-tier entries in Figure 8. **Medium tier**: Secretory cell (0.719), Excitatory neuron (0.697), Oligodendrocyte precursor (0.691), Epithelial cell METTL12+ (0.603), Innate lymphoid cell (0.508), CD8+ T cell (0.422), CD4+ T cell FOSB+ (0.406), CD8+ T cell KLRG1+ (0.405). **Low tier**: includes Macrophage Metallothionein+ (0.245), Fibroblast SFRP2+ SLPI+ (0.115), and three zero-accuracy types (Adrenal cortical cell, Schwann cell, Fibroblast). The contraction of the High tier from eight to two entries foreshadows the cross-task difficulty gradient.

**Figure 10.**
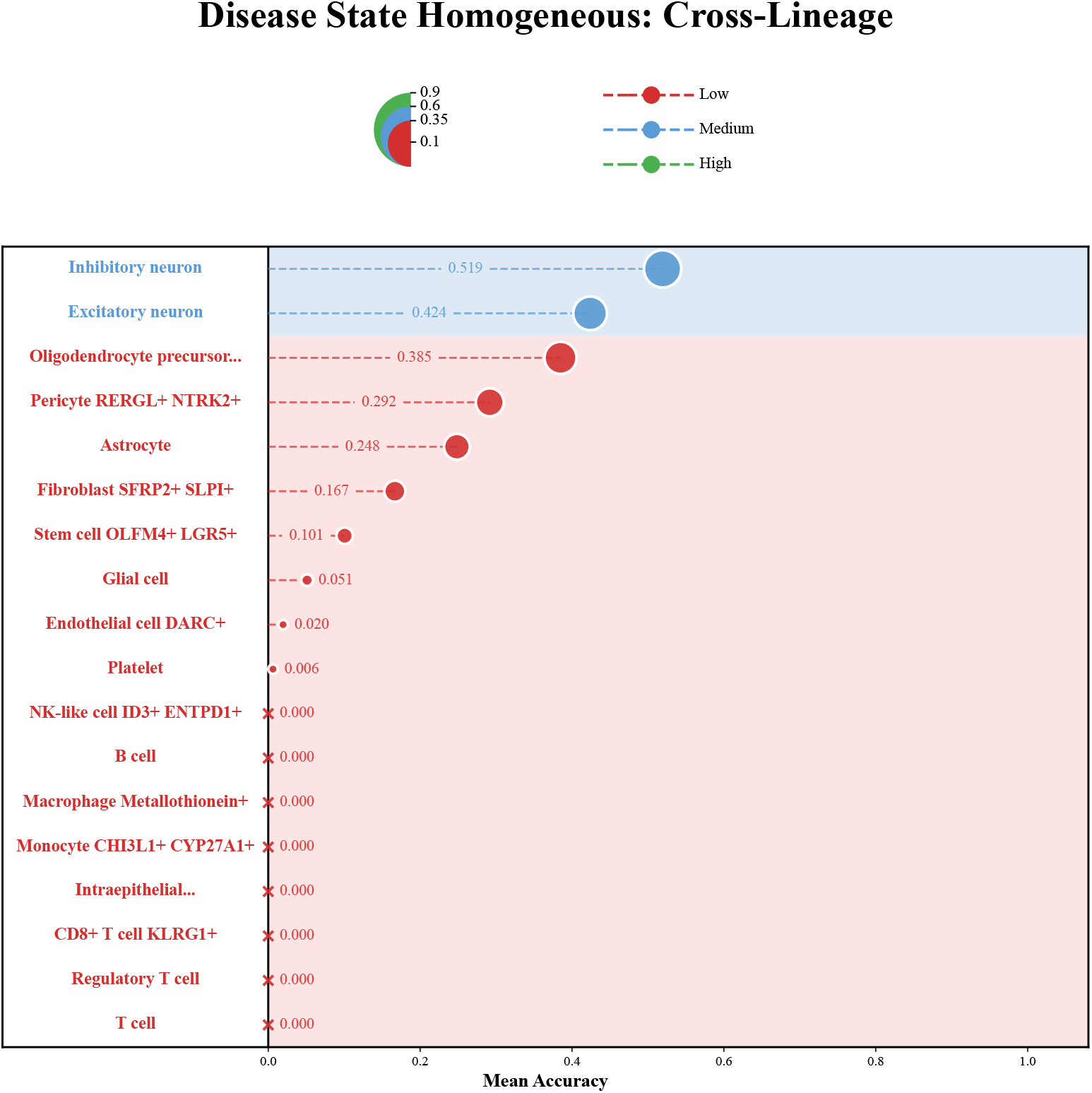
Per-cell-type accuracy distribution — disease state classification. Lollipop chart of per-cell-type accuracy on the homogeneous disease-state task (the converged GSPO disease-state policy; offline evaluation judge threshold 0.85—stricter than the 0.75 used for cell and tissue type, reflecting more demanding evaluation for disease state). **No cell type reaches the High tier** (≥0.8), completing the difficulty gradient cell type (8 High) → tissue type (2 High) → disease state (0 High). **Medium tier**: only Inhibitory neuron (0.519) and Excitatory neuron (0.424). **Low tier**: Oligodendrocyte precursor (0.385), Pericyte RERGL+ NTRK2+ (0.292), Astrocyte (0.248), Fibroblast SFRP2+ SLPI+ (0.167), Stem cell OLFM4+ LGR5+ (0.101), Glial cell (0.051), Endothelial cell DARC+ (0.020), Platelet (0.006), plus eight zero-accuracy types (NK-like cell, B cell, Macrophage Metallothionein+, Monocyte CHI3L1+, Intraepithelial lymphocyte, CD8+ T cell KLRG1+, Regulatory T cell, and T cell). The absence of any High-tier entry and the prevalence of zero-accuracy immune cell types confirm that disease-state prediction is the hardest of the three homogeneous-population tasks.

**Figure 11.**
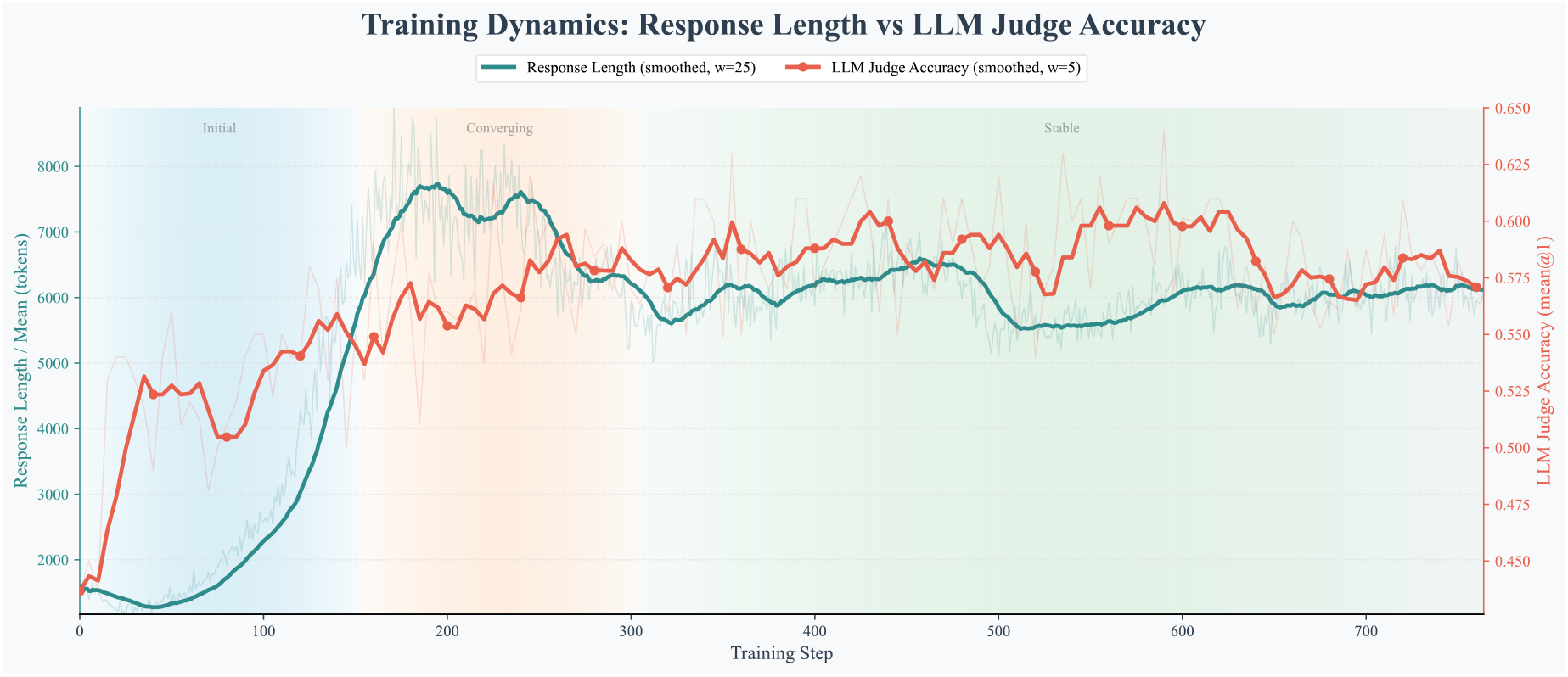
Tissue type — response length vs. accuracy. Companion to Figure 6a for homogeneous tissue-type classification (the converged GSPO tissue policy). Three training phases are annotated: Initial (~ 0–150 steps), Converging ( ~150–300), and Stable ( ~300–770). Response length (teal, rolling window of 25 steps) starts ~1,500 tokens, peaks at ~7,700 around steps 180–200, and settles to ~6,000– 6,200 in the Stable phase. LLM judge accuracy (coral, rolling window of 5 steps) starts ~0.44, climbs to ~0.62 near step 190, then hovers at 0.55–0.61 through the Stable phase with local peaks ~0.61 near steps 430–440 and 570–620. Both curves exhibit a much smaller dynamic range than the cell-type figure (Figure 6a), suggesting that the tissue-type task reaches a training plateau earlier and with less dramatic length compression.

**Figure 12.**
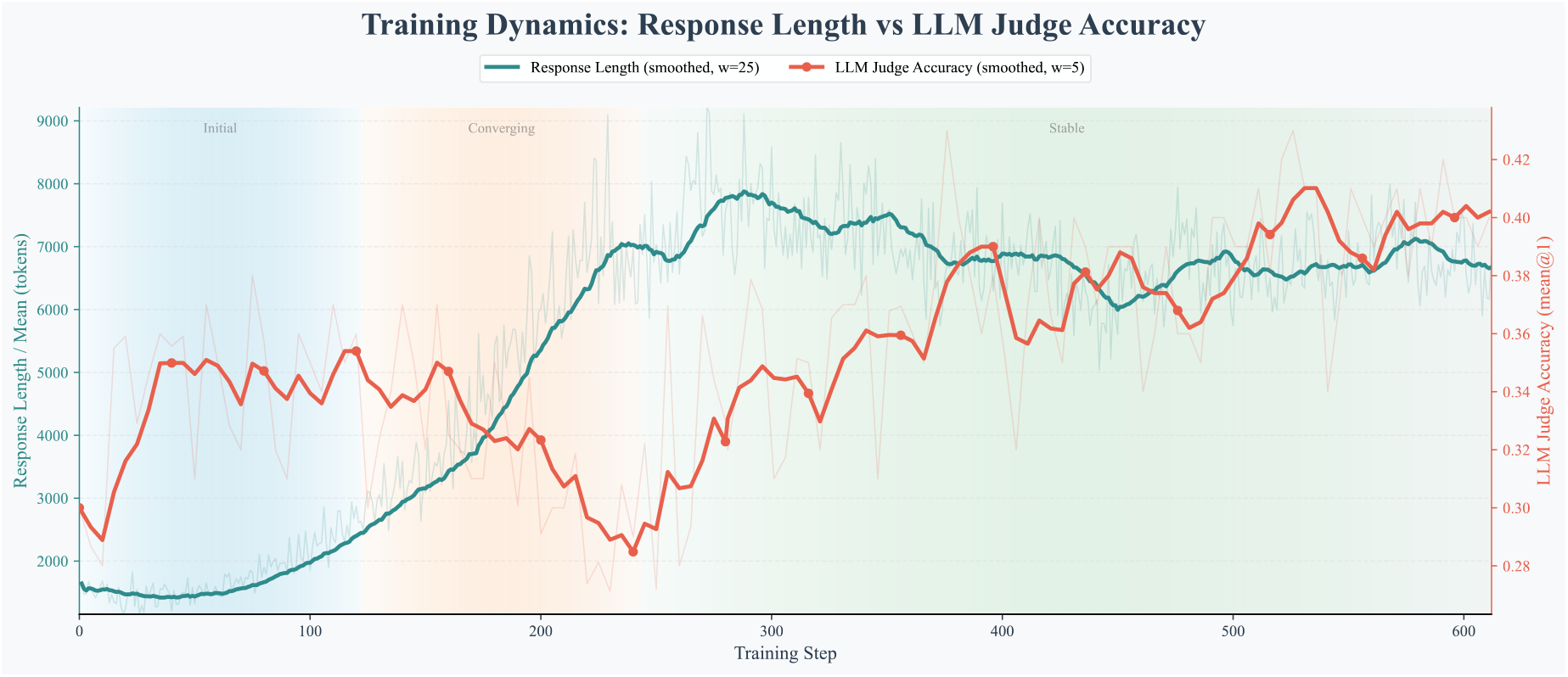
Disease state — response length vs. accuracy. Companion to Figure 6a for homogeneous disease-state classification (the converged GSPO disease-state policy). Three phases: Initial (~0–120 steps), Converging ( ~120–260), Stable ( ~260–610). Response length (teal) starts ~1,500–1,600 tokens, peaks at ~7,800–7,900 around step 290, then fluctuates between 6,500–7,100 through the Stable phase. Accuracy (coral) occupies the narrowest range of the three tasks (0.28–0.42): it starts ~0.30, rises to ~0.35 by step 40, drops to a trough of ~0.285–0.29 around step 240 in the Converging phase, then rebounds unevenly through Stable with oscillations (~0.39–0.40 at step 420, dip to ~0.36 at step 480, peak ~0.41 at step 535), ending near ~0.40. Disease state shows the weakest length-compression effect and the noisiest accuracy trajectory, consistent with it being the hardest task.

**Figure 13.**
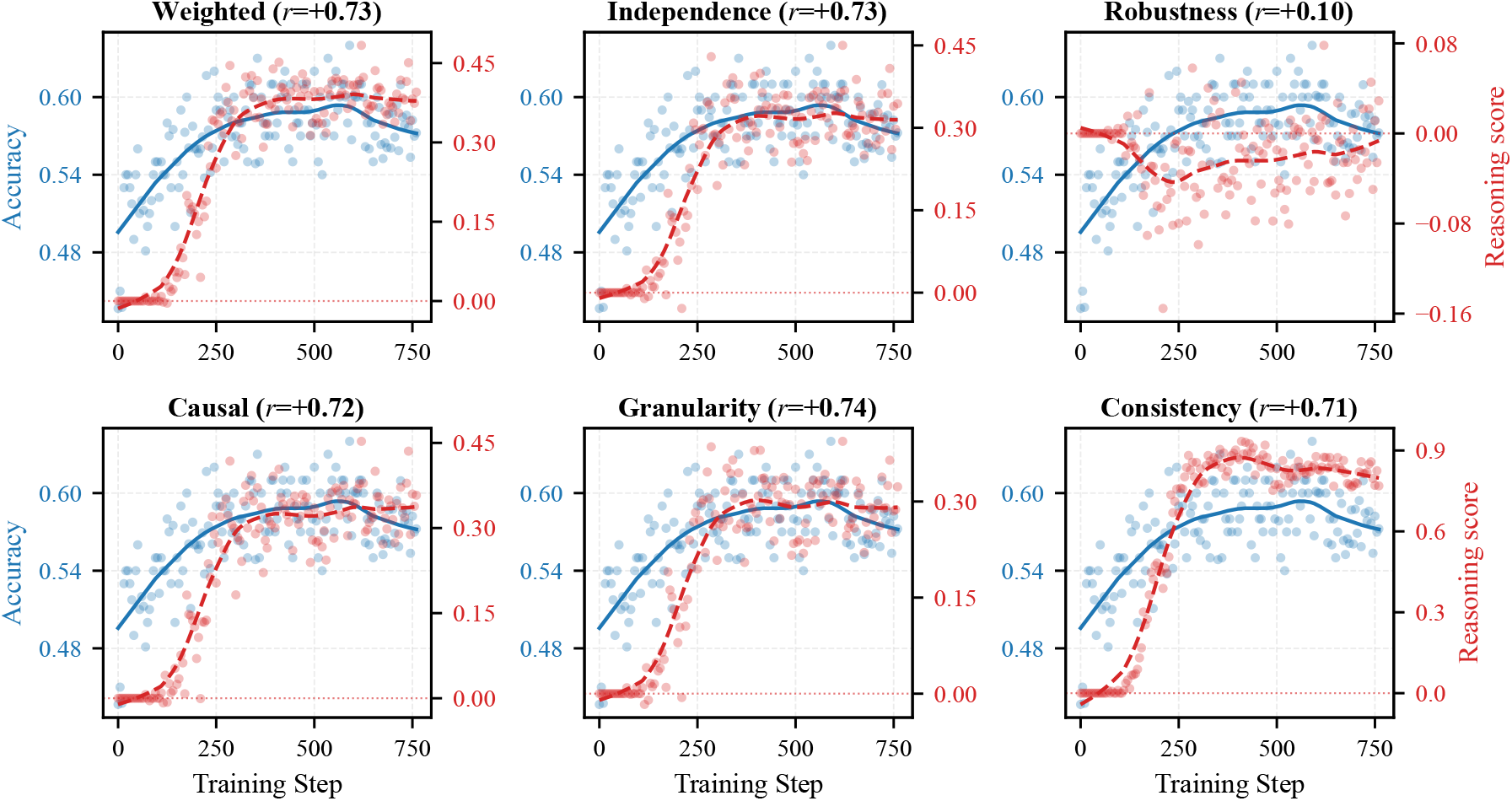
Tissue type — accuracy vs. reasoning quality. Companion to Figure 6b for homogeneous tissue-type classification (the converged GSPO tissue policy, ~ 750 training steps). LLM judge accuracy (blue, left axis, range 0.450–0.650) rises from ~ 0.495 to a plateau near ~ 0.59–0.60 from step ~300, with a slight terminal downturn. Per-dimension Pearson correlations: Granularity (*r* = +0.74), Reasoning Quality Weighted (*r* = +0.73), Independence (*r* = +0.73), Causal (*r* = +0.72), Consistency (*r* = +0.71), and Robustness (*r* = +0.10). Most reasoning curves (red, right axis) rise sharply between steps ~100– 300 and then plateau, mirroring the accuracy trajectory. All correlations are noticeably weaker than the cell-type range [0.60, 0.92]; Robustness is essentially flat (*r* = +0.10), foreshadowing its collapse to negative values on disease state (Figure 14).

**Figure 14.**
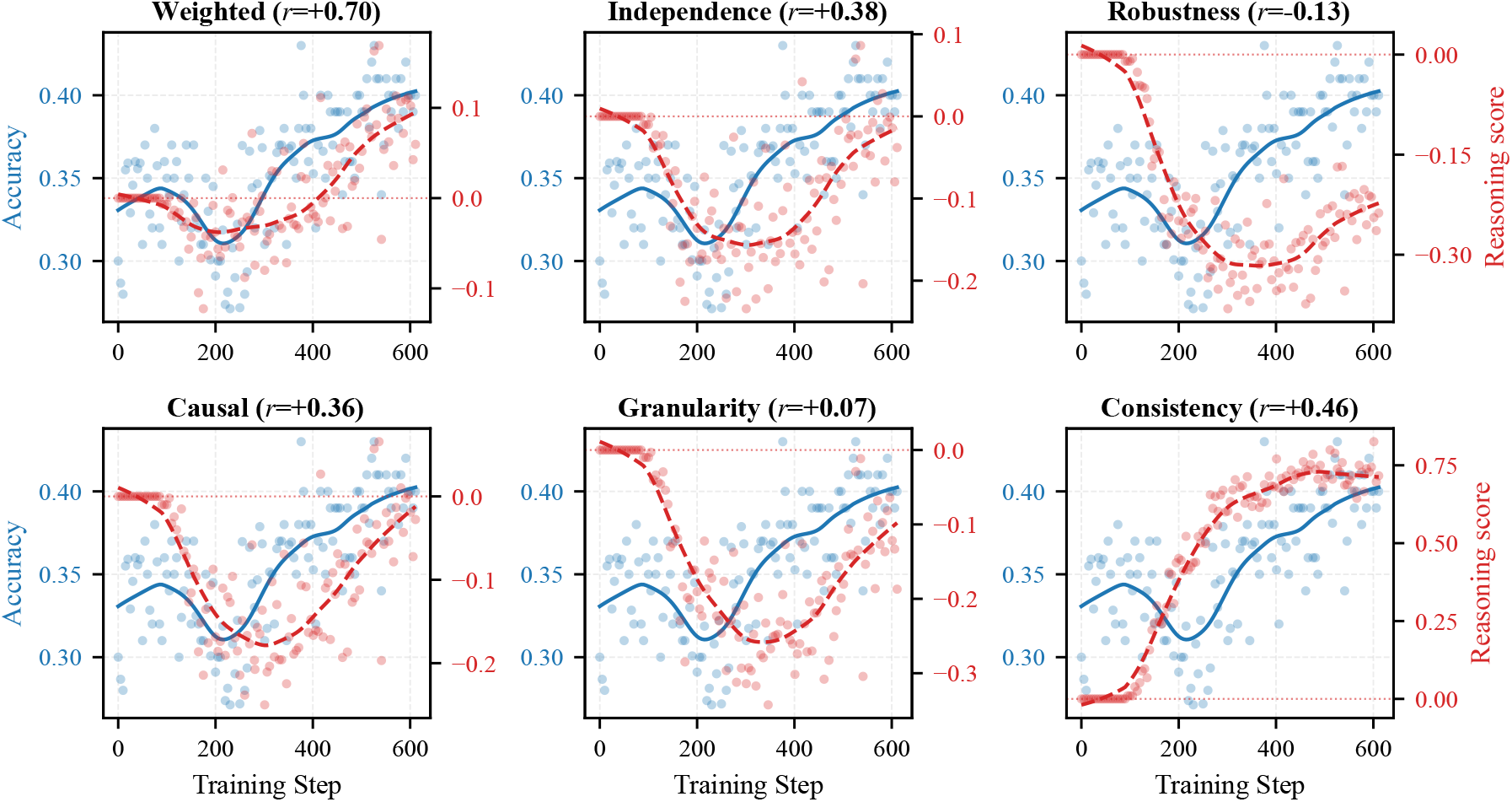
Disease state — accuracy vs. reasoning quality. Companion to Figure 6b for homogeneous disease-state classification (the converged GSPO disease-state policy, ~600 training steps). LLM judge accuracy (blue, left axis, range 0.28–0.42) follows a U-shaped recovery: starts ~0.33, dips to ~0.31 around steps 200–250, then rises to ~0.40 by step 600. Per-dimension Pearson correlations: Reasoning Quality Weighted (*r* = +0.70), Consistency (*r* = +0.46), Independence (*r* = +0.38), Causal (*r* = +0.36), Granularity (*r* = +0.07), and Robustness (*r* = − 0.13). Two dimensions collapse on this hardest task: **Robustness** turns *negative* (*r* = −0.13; right-axis range − 0.40 to 0.00, monotonic decline then partial recovery), the only negative correlation among the 18 (task, dimension) pairs. **Granularity** drops to near zero (*r* = +0.07; right-axis range − 0.35 to 0.00), down from *r* = +0.92 on cell type. This forms the cross-task correlation gradient: Cell Type (strong, *r* = 0.60–0.92) > Tissue Type (moderate, *r* = 0.10–0.74) > Disease State (weak, *r* = − 0.13 to +0.70), providing the strongest evidence that the reward signal loses traction on hard tasks and motivating task-specific reward tuning as future work.

**Table 8:**
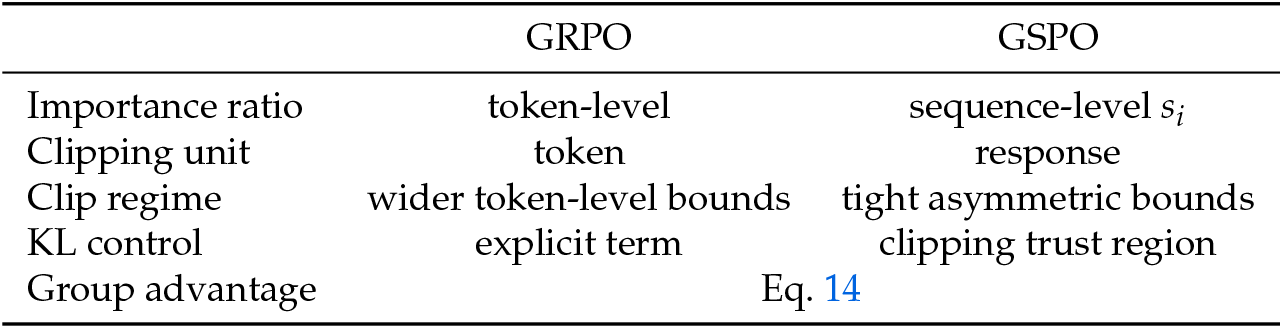
GRPO [50] and GSPO [47] share the group-relative advantage estimator but apply importance weighting and clipping at different granularities.

**Table 9:** Reward-side answer judge scoring categories. The training-time reward-side LLM judge classifies each prediction into one of seven categories reflecting positions on the Cell Ontology graph. These scores enter the training-time reward signal only; all accuracy numbers reported in the main text are produced by the independent evaluation protocol of §4.4. Score ranges are continuous within each category.

| Category | Score Range | Ontological Interpretation |
| --- | --- | --- |
| Exact Match | +0.9 to +1.0 | Prediction matches ground truth exactly |
| More Specific | +0.4 to +0.85 | Valid sub-type in the ontology hierarchy |
| Less Specific | −0.05 to 0.0 | Valid ancestor; correct but imprecise |
| Wrong Branch | −0.3 to −0.6 | Same lineage, different branch |
| Vague / Evasive | −0.1 to −0.3 | Non-committal or hedging response |
| Wrong Lineage | −0.7 to −0.9 | Different cell lineage entirely |
| Unrelated | −0.95 to −1.0 | No meaningful biological relationship |

**Table 10:** Reasoning evaluation dimensions. Five dimensions carry non-uniform weights and are aggregated into *s*_reason_ (Eq. 7). A sixth dimension is logged for monitoring but excluded from the reward to avoid double-counting with the answer judge.

| Dimension | Weight | Definition |
| --- | --- | --- |
| Independence ($d_{\text{ind}}$) | 0.30 | Whether conclusions derive from gene expression evidence rather than restating the query or relying on prior knowledge without grounding |
| Counterfactual Robustness ($d_{\text{rob}}$) | 0.25 | Whether reasoning holds under different tissue or disease contexts, ensuring cell-intrinsic rather than dataset-specific features |
| Causal Sufficiency ($d_{\text{caus}}$) | 0.20 | Logical necessity between cited evidence and the stated conclusion; penalizes leaps of inference |
| Granularity Discipline ($d_{\text{gran}}$) | 0.15 | Appropriateness of specificity relative to available evidence; penalizes over- or under-specific claims |
| Consistency ($d_{\text{cons}}$) | 0.10 | Mutual agreement between cited gene evidence and the final cell-type answer |
| Label Semantic Match ($d_{\text{sem}}$) | — | Alignment between the reasoning chain’s implied cell identity and the ground-truth label; logged for monitoring, excluded from reward to avoid double-counting with $R_{\text{answer}}$ |

**Table 11:** RVQ tokenizer architecture configuration. All values correspond to the released model. The encoder is a two-layer MLP with hidden widths 1024 and 512 mapping *G* → 128; each linear layer is followed by batch normalisation and a leaky ReLU activation. The decoder mirrors this architecture (128 → 512 → 1024 → *G*).

| Parameter | Description | Value |
| --- | --- | --- |
| Input dimensionality ($G$) | Number of measured genes | 36,601 |
| Hidden dimensionalities | Encoder/decoder hidden layer sizes | [1024, 512] |
| Latent dimensionality ($D$) | Latent vector dimensionality | 128 |
| Number of codebooks ($N_c$) | Number of RVQ codebook levels | 8 |
| Codebook size ($C$) | Entries per codebook | 32 |
| Quantization weight ($\lambda_q$) | Quantization loss weight | 0.1 |
| Total RVQ vocabulary tokens ($N_c \times C$) | | 256 |
| Information rate per cell ($N_c \cdot \log_2 C$) | | 40 bits |
| Encoder parameters | | $\sim 38.1$ M |
| Decoder parameters | | $\sim 38.1$ M |
| Codebook parameters ($N_c \times C \times D$) | | 32,768 |
| **Total parameters** |  | **$\sim 76.2$ M** |

**Table 12:** Per-codebook utilization and label association. Each entry is training/holdout (T/H); usage gives active codes out of 32, entropy uses natural logarithms, and NMI is arithmetic-normalized. Cell-type mapping coverage is 98.5%/100% while lineage coverage is 93.1%/40.4%, so holdout lineage NMI is auxiliary evidence.

| CB | Usage T/H | Entropy T/H | PPL T/H | Cell NMI T/H | Lineage NMI T/H | Dataset NMI T/H |
| --- | --- | --- | --- | --- | --- | --- |
| 0 | 28/16 | 1.8364/1.6299 | 6.27/5.10 | 0.4458/0.4461 | 0.5469/0.5286 | 0.4208/0.5128 |
| 1 | 32/27 | 2.9116/2.7192 | 18.39/15.17 | 0.4621/0.4654 | 0.2390/0.3413 | 0.3391/0.3675 |
| 2 | 32/31 | 2.9638/2.6747 | 19.37/14.51 | 0.3153/0.2821 | 0.1575/0.1735 | 0.3312/0.2711 |
| 3 | 32/31 | 3.0947/3.0575 | 22.08/21.27 | 0.2538/0.2944 | 0.0752/0.1480 | 0.2165/0.2442 |
| 4 | 32/30 | 3.0659/2.8740 | 21.45/17.71 | 0.2140/0.2674 | 0.0741/0.1691 | 0.1961/0.2168 |
| 5 | 32/32 | 3.0630/2.9930 | 21.39/19.95 | 0.1624/0.2265 | 0.0524/0.1243 | 0.1447/0.1849 |
| 6 | 32/32 | 3.2141/3.1804 | 24.88/24.06 | 0.1901/0.2451 | 0.0756/0.1289 | 0.1809/0.2078 |
| 7 | 32/31 | 3.1427/3.1288 | 23.17/22.85 | 0.1378/0.2029 | 0.0577/0.1019 | 0.1168/0.1820 |

**Table 13:** Prefix-*K* accumulation for the deployed RVQ tokenizer. Brackets give 95% dataset-bootstrap intervals. *K* = 0 passes a zero latent to the decoder rather than representing a zero-expression cell; prefix decoding is out of distribution because the decoder was trained on full eight-codebook sums.

| $K$ | Residual energy ↓ | Recon. MSE ↓ | Per-cell Pearson ↑ | Cosine ↑ |
| --- | --- | --- | --- | --- |
| 0 | 1.0000 [1.0000, 1.0000] | 0.2273 [0.2104, 0.2461] | 0.4419 [0.3876, 0.4909] | 0.4713 [0.4279, 0.5116] |
| 1 | 0.2107 [0.1618, 0.2643] | 0.0886 [0.0775, 0.1011] | 0.5751 [0.5261, 0.6174] | 0.5884 [0.5472, 0.6237] |
| 2 | 0.1651 [0.1204, 0.2152] | 0.0881 [0.0773, 0.1005] | 0.6031 [0.5529, 0.6443] | 0.6151 [0.5715, 0.6505] |
| 4 | 0.1303 [0.0914, 0.1742] | 0.0884 [0.0773, 0.1008] | 0.6228 [0.5701, 0.6668] | 0.6337 [0.5876, 0.6722] |
| 8 | 0.0953 [0.0646, 0.1302] | 0.0894 [0.0774, 0.1025] | 0.6361 [0.5827, 0.6803] | 0.6466 [0.5996, 0.6852] |

**Table 14:** Paired prompt length and single-request prefill efficiency. Token counts are medians over every record in each stratum. Prefill uses the same Qwen3-4B GSPO checkpoint and a vLLM 0.18.0 backend on one H100 80GB GPU, with pretokenized inputs, prefix caching disabled, concurrency and batch size one, one generated token, three deterministic pairs per stratum, and three repetitions per arm; it excludes host tokenization and long-response decoding

| Scope | Records | Cells | Text tok. | RVQ tok. | Reduction / Prefill |
| --- | --- | --- | --- | --- | --- |
| CTA | 3,840 | 1 | 780 | 343 | 55.9% / $1.02\times$ |
| TI Natural, $N = 8$ | 201 | 8 | 3,963 | 416 | 89.5% / $3.31\times$ |
| TI Natural, $N = 16$ | 201 | 16 | 7,625 | 521 | 93.2% / $6.91\times$ |
| TI Natural, $N = 32$ | 201 | 32 | 14,940 | 729 | 95.1% / $14.53\times$ |
| TI Natural, $N = 64$ | 201 | 64 | 29,602 | 1,145 | 96.1% / $39.95\times$ |
| DSA Natural, $N = 8$ | 30 | 8 | 3,948 | 390 | 90.1% / $3.33\times$ |
| DSA Natural, $N = 16$ | 30 | 16 | 7,624.5 | 495.5 | 93.5% / $6.54\times$ |
| DSA Natural, $N = 32$ | 30 | 32 | 14,944 | 704 | 95.3% / $15.34\times$ |
| DSA Natural, $N = 64$ | 30 | 64 | 29,603.5 | 1,121 | 96.2% / $38.51\times$ |
| PCA cross-tissue | 196 | 64 | 29,246.5 | 1,043 | 96.4% / $39.98\times$ |
| PCA cross-disease | 105 | 64 | 29,737 | 1,044 | 96.5% / $41.02\times$ |

**Table 15:** Cross-method tokenizer comparison grouped by bit budget. Bits = *N*_*c*_ log_2_ *C*; *Usage* is codebook utilization; ARI/AMI/NMI are *K*-means agreement with broad cell-type labels. This held-out measurement differs from the configuration study of Table 7, so the two are not directly comparable. Bold marks the best value within a budget group containing multiple models; the single 96-bit row is contextual.

| Tokenizer | Steps | Bits | Vocab | Usage % | MSE ↓ | PCC<sub>c</sub> ↑ | PCC<sub>g</sub> ↑ | ARI ↑ | AMI ↑ | NMI ↑ |
| --- | --- | --- | --- | --- | --- | --- | --- | --- | --- | --- |
| *40-bit matched budget* |  |  |  |  |  |  |  |  |  |  |
| RVQ (8×32)* | 25,500 | 40 | 256 | **85.94** | 0.0973 | **0.680** | **0.179** | **0.562** | **0.698** | **0.698** |
| PQ (8×32) | 15,000 | 40 | 256 | 77.34 | **0.0921** | 0.678 | 0.171 | 0.499 | 0.639 | 0.639 |
| *96-bit context* |  |  |  |  |  |  |  |  |  |  |
| RVQ (16×64) | 15,000 | 96 | 1024 | 22.36 | 0.0878 | 0.678 | 0.171 | 0.517 | 0.680 | 0.681 |
| *256-bit context* |  |  |  |  |  |  |  |  |  |  |
| RVQ (32×256) | 15,000 | 256 | 8192 | 17.99 | 0.0881 | 0.682 | 0.174 | 0.495 | 0.663 | 0.664 |
| PQ (32×256) | 15,000 | 256 | 8192 | **39.69** | 0.0858 | **0.697** | **0.193** | 0.493 | 0.666 | 0.667 |
| CellTok-style reimpl. | 15,000 | 256 | 8192 | 4.03 | **0.0831** | 0.691 | 0.164 | **0.540** | **0.670** | **0.671** |

## A Misclassification Flow

This appendix retains the alluvial decomposition of cell-type misclassification flows referenced from the main text.

## B Per-Cell-Type Accuracy Breakdown

These lollipop charts expose the cross-task difficulty gradient under the task-specific population evaluation protocol (§4.4): cell type has multiple perfect-performance entries, tissue type has fewer, and disease state has none in the High tier. Each chart classifies cell types into three tiers—High (≥0.8, green), Medium (0.4–0.8, blue), and Low (< 0.4, red)—with bubble area proportional to accuracy and a small fixed minimum size to keep low-accuracy types visible; zero-accuracy types are marked with “×”. Reporting grain: the post-training offline evaluation judge emits a per-cell score in [0, 1]; for these per-cell-type breakdowns we apply the task-dependent threshold at the cell grain (*score*≥*task-specific threshold* →1, else 0) and then average these per-cell binary scores within each cell-type bucket to obtain the value plotted here—a cell-level grain distinct from the deterministic per-record scoring of Table 5. Thresholds are task-specific (cell type: 0.75; tissue type: 0.75; disease state: 0.85) rather than globally uniform, and are noted in each caption.

## C Cross-Task Reward Dynamics

These figures extend the main-text length and reasoning-quality dynamics (Figure 6) to tissue and disease state classification. Together with the main-text cell-type figures, they reveal a systematic cross-task gradient: reasoning-quality correlations weaken from cell type (*r* ∈ [0.60, 0.92]) through tissue type (*r* ∈ [0.10, 0.74]) to disease state (*r* ∈ [ −0.13, 0.70]). Response-length and accuracy curves are smoothed with rolling windows of width 25 and 5 respectively; reasoning-quality trends are smoothed with LOWESS (smoothing fraction 0.3), and per-dimension Pearson correlations *r* are reported.

## D Sequence-Level Policy Optimization

Biological predictions and their rationales are evaluated as complete responses: the validity of a biological conclusion depends on the answer and the evidence chain taken together. Group Sequence Policy Optimization (GSPO) [47] matches this granularity by assigning one importance ratio to each sampled sequence, while retaining the group-relative comparison used by Group Relative Policy Optimization (GRPO) [50].

### Group-normalized advantages

For each prompt *q*, the policy samples *N*_*g*_ candidate outputs 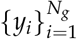 and standardizes their rewards within the group:

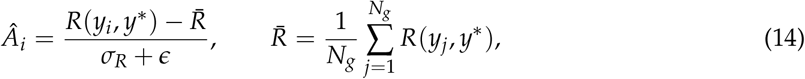

where *R*(*y*_*i*_, *y*^∗^) is the clipped composite reward defined in Eqs. 6–9. The within-group baseline adapts the update scale to the reward distribution of the current prompt, which is useful when task routes differ in difficulty and score range.

### Sequence-level importance ratio

GSPO summarizes the per-token likelihood changes of a response by their geometric mean:

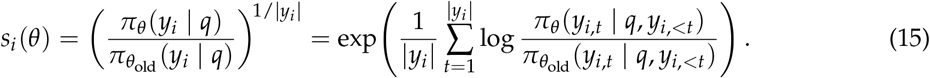

Length normalization makes responses of different sizes comparable, while the single ratio keeps the trust-region decision aligned with the response-level reward. Table 8 summarizes the distinction from token-level GRPO.

### Asymmetric clipped surrogate

The policy minimizes the response-level clipped surrogate

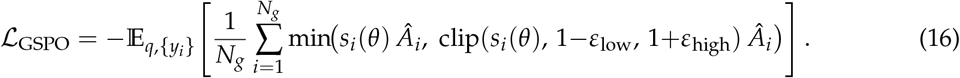

The tight asymmetric bounds provide the trust region for the sequence-level ratio, so the objective does not add a separate per-token KL penalty. Section 4.1 reports the adopted bounds and optimization configuration.

## E Reward System Design Details

This section provides the implementation specifications for the reward system components summarized in the main text (Section 3.4.1 and Section 3.4.2).

### Pre-gates

Before the multiplicative formula (Eq. 6), two sequential binary gates can zero out 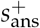:

- **Gate A**^′^ **(format compliance)**. Outputs that omit the dedicated reasoning span (the paired `<think>` and `</think>` delimiters), fall below task-specific minimum reasoning lengths, or yield an empty answer have 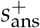 zeroed, forcing the negative path.
- **Gate A**^′′^ **(answer output contract)**. If the answer violates structural constraints—marker stuffing (specificity score > 2), template regurgitation (placeholder patterns such as `<answer>`), or answer repetition (unique token ratio < 0.3)—then 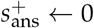.

Both gates block only the positive path; 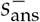 flows through unchanged, maintaining the penalty gradient for wrong predictions. Preserving the negative path regardless of format validity prevents the model from escaping penalty by deliberately producing malformed output.

### Anti-reward-hacking defenses

Six defense layers, added iteratively as reward-hacking modes were observed during training, guard against reward exploitation. Binary gates (which short-circuit computation) precede continuous penalties (which modulate magnitude):

- **System message isolation**. Judge instructions are placed in the system role and untrusted model output is sandboxed in the user role with structured escaping, preventing prompt-injection against the judge.
- **Adversarial input detection**. An adversarial-input filter spans five attack categories (score manipulation, dimension-name injection, system-override attempts, fabricated JSON, and multitag abuse); samples matching two or more patterns receive an immediate penalty of − 0.8 before any judge evaluation.
- **Format gate (A**^′^). Blocks positive reward path for structurally invalid outputs (see the *Pre-gates* paragraph above).
- **Answer output contract (A**^′′^). Six sequential checks—template regurgitation (−0.5), answer overlong (−0.2 gradual), marker stuffing (−0.5 gradual, threshold exceeds *w*_base_ by design), answer repetition (−0.15 gradual), truncation without EOS (−0.3), and answer leakage into reasoning (−0.15).
- **Repetition detection**. A 4-gram overlap penalty applies independently to the reasoning trace (cap −0.2) and to the full output (cap −0.5), taking the more severe of the two.
- **Length penalty**. A sub-linear length penalty activates above a task-specific character budget and saturates at a hard cap of roughly twice that budget.

A key design principle: the marker stuffing penalty ( − 0.5) intentionally exceeds *w*_base_ = 0.3, ensuring that even a semantically correct answer with marker stuffing produces a net negative reward.

### Answer judge scoring categories

Table 9 lists the seven graded categories used by the *reward-side* ontology-aware answer judge, with their score ranges and ontological interpretations. This rubric, and all multi-sample aggregation described in this appendix, are properties of the training-time reward system; they are not used by the post-training offline evaluation judge that produces the main-text accuracy numbers (§4.4), which operates under a different semantic-relation schema and without reward-side multi-sample aggregation.

During training-time reward computation, to mitigate judge variance each prompt is evaluated *N* = 3 times by the reward-side answer judge. Within these three samples, the scalar score is aggregated by the *mean* and feeds the reward signal, while the *median* is used solely for the internal reasoning-gate decision (whether to invoke the reasoning judge); when the seven-category vote is tied, the reward system applies conservative tie-breaking toward the lower-scoring category. This aggregation is not applied by the post-training offline evaluation judge described in §4.4.

### Reasoning dimension definitions

Table 10 provides full definitions for the five weighted reasoning dimensions and the monitored-only sixth dimension.

The non-uniform weighting encodes domain expertise: independence and counterfactual robustness— the dimensions most diagnostic of genuine biological reasoning versus superficial pattern matching— receive the largest weights.

### Population aggregation

For population-level tasks (multi-cell inputs), the reward system decomposes evaluation to the per-cell level. Each of *N* cells receives an independent answer judge evaluation, and the scalar signals are aggregated:

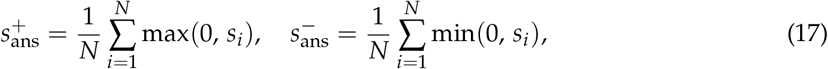

where *s*_*i*_ is the normalized answer score for cell *i*. The aggregated signals feed the same multiplicative gate (Eq. 6). This decomposition prevents a failure mode where the model achieves a high population-level score by correctly predicting the majority cell type while ignoring minority populations. Additional population-specific penalties enforce consistency: a homogeneous population with inconsistent predictions incurs *P*_consistency_ = −0.3; missing cells receive −0.2 each and extraneous cells −0.1 each.

### Fact verification infrastructure

Each reasoning chain *y* is decomposed into typed atomic claims *C*={*c*_1_, …, *c*_*K*_}. An untrusted extractor proposes minimal semantic units together with their sentence anchors; a deterministic compiler alone fixes character spans, cell scopes, and gene identities. Four fail-closed filters govern admission: verbatim reproduction of the source span up to unambiguous span repair, a type and scope drawn from the current rollout’s allowlist, content-keyed deduplication that never merges evidence across cells, and explicit exposure of inter-claim dependencies without credit flowing through them.

Evidence combines marker context (CellMarker, scMarkerAgent, Human Protein Atlas), regulon and program activity (Hallmark, PROGENy, CollecTRI, DoRothEA), frozen ontology relations (Cell Ontology, UBERON, DOID, MONDO, PATO), and the cell’s own RVQ-decoded expression profile. Each admitted claim is resolved along three orthogonal axes—truth, grounding, and overclaim—to one of Supported, Contradicted, or Uncertain. Missing catalog entries, non-human samples, and genes absent from a rank-truncated profile resolve to Uncertain; mixed-lineage, dissociation, and cell-cycle guards may withhold evidence but cannot create a contradiction. A two-stage cache (exact match followed by embedding-based match) amortizes verification across recurring claims, achieving a hit rate above 80% in our runs.

Evidence is served only from immutable, content-hashed sidecars, and a recursive guard strips the verification context of any structure carrying target-, teacher-, or reference-keyed fields. A lookup miss resolves to Uncertain without reopening a source expression matrix; that outcome carries no penalty and does not proxy biological absence.

### Response-level factuality diagnostics

Each response is assigned to one of four mutually exclusive categories under a fixed priority: Missing/Refusal/Unscorable, Contradiction, Obligation-Only, or Safe-Auditable. The first three form the primary-unsafe set, and Unscorable takes precedence so that a response which could not be audited does not inflate the contradiction count. Writing *n*_S_, *n*_C_, and *n*_U_ for the supported, contradicted, and uncertain claim counts, with resolved count *r* = *n*_S_ + *n*_C_ and adjudicated count *a* = *r* + *n*_U_, the resolved hallucination rate is *H*_res_ = *n*_C_/*r* and coverage is *r*/*a*; the interval [ *n*_C_/*a*, (*n*_C_ + *n*_U_)/*a*] brackets the rate by treating every uncertain claim first as correct and then as wrong. These diagnostics are reported separately from the bounded adjustment of Eq. 8 and never modulate it.

### Reward hyperparameters

The multiplicative gate (Eq. 6) uses fixed weights *w*_base_ = 0.3, *w*_reason_ = 0.5, and *w*_neg_ = 1.0, with a reasoning judge gate threshold *τ*_gate_ = 0.3. The bounded contradiction adjustment (Eq. 8) uses the active V1 policy *P*_max_ = 0.4, *λ*_any_ = 0.3, *λ*_count_ = 0.1, and *K*_*c*_ = 1, so any nonzero eligible contradiction count yields *P*_fact_ = −0.4.

## F RVQ Tokenizer Architecture, Training, and Vocabulary Expansion

This supplementary consolidates the implementation details abstracted away in §3.2: the deployed architecture configuration, the full training objective, the augmentation curriculum, the embedding-initialization procedure, and the vocabulary-expansion mechanics.

### Architecture configuration

Table 11 lists the hyperparameters of the deployed RVQ autoencoder, matching the values released alongside the public checkpoint.

The compression cascade 36,601 → 1024 (~35.7×) → 512 (2×) → 128 (4×) follows a smooth geometric reduction, consistent with the encoder backbones of SoundStream [38] and VQ-VAE-2 [37]. The encoder hidden widths match those of CellTok [19] (5000 → 1024 → 512 → 256), providing direct domain precedent for this configuration.

### Training objective

The tokenizer is trained end-to-end with a composite loss:

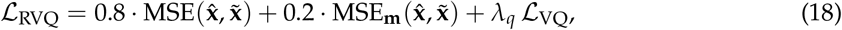

where 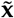 is the post-scaling, pre-dropout target, **m** marks the coordinates zeroed by gene dropout, and 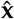 is the reconstruction obtained from the masked input 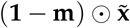. A single forward pass produces 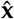: MSE averages over all *G* coordinates and MSE_**m**_ over those with *m*_*g*_ = 1, so the two terms differ in which coordinates they score rather than in their target. We set *λ*_*q*_ = 0.1. The quantization loss comprises commitment, embedding, and usage regularizer terms:

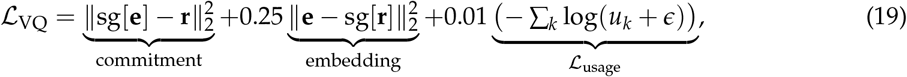

where sg[·] is the stop-gradient operator and *u*_*k*_ is the fraction of unique codes used in codebook *k*. The usage regularizer prevents codebook collapse by penalizing low usage.

### Augmentation curriculum

An adaptive curriculum applies Gaussian noise and gene dropout during early training, with both augmentation magnitudes decayed exponentially after epoch 10. The masked-coordinate term in Eq. 18 enforces robustness to sequencing sparsity: the decoder must recover the zeroed entries of 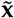 from an encoder input in which they are absent.

### Vocabulary expansion and embedding initialization

The 256 RVQ code tokens produced by Eq. 2 are appended to the text vocabulary of the base LLM, together with five structural delimiter tokens (`<|singlecell_start|>, <|singlecell_end|>, <|singlecell_interval|>, <|multicell_start|>,` and `<|multicell_end|>`) that bracket cells at the per-cell and population levels. For the deployed base model (Qwen3-4B-Thinking-2507), the tokenizer holds 151,669 entries before expansion; this is the quantity the offset is computed from, and it is distinct from the 151,936-row embedding matrix the model configures, which is padded for alignment. Adding the five delimiters gives *V*_text_ = 151,674, and the RVQ tokens then occupy the contiguous block [*V*_text_, *V*_text_ + *N*_*c*_*C*) = [151,674, 151,930), adding exactly 256 new entries. Six reserved slots pad the total back to 151,936, a multiple of eight, so the expanded tokenizer matches the configured embedding matrix and no resize is required. Embeddings for the new tokens are initialized from *N* (0, *σ*^2^), matching the scale of the pretrained text embeddings, with a diversity verification step that resamples any pair whose cosine similarity exceeds 0.5. This procedure keeps the 256 newly introduced tokens well separated at initialization without disrupting the existing text embedding distribution.

### Codebook utilization and label association

Table 12 audits the deployed eight-codebook representation on all 56 training shards and 13 holdout shards. The first codebook is markedly more concentrated, whereas later codebooks use nearly the full inventory at higher perplexity. Cell-type association peaks at CB1 and lineage association at CB0; the decline thereafter is compatible with progressively finer residual allocation but does not by itself identify an ordered biological hierarchy. The sizable dataset NMI at CB0 (0.4208/0.5128) rules out the stronger reading that early codebooks encode biological lineage alone.

### Prefix-*K* residual refinement

Using the released tokenizer checkpoint, we deterministically sampled 512 cells from each of the 13 holdout datasets and re-encoded the raw profiles in float32. D102 is retained only as a diagnostic because of legacy checkpoint-selection exposure, so Table 13 reports an unweighted macro over the remaining 12 datasets with 95% intervals from 10,000 dataset-level bootstrap replicates. All 53,248 frozen code positions were reproduced exactly, and the *K* = 8 latent and reconstruction matched the native full-quantization path.

The first codebook removes 78.9% of normalized latent residual energy and all eight remove 90.5%; dataset-level mean residual energy decreases at every reported prefix for all 13 datasets. Pearson and cosine similarity also rise with *K*. Reconstruction MSE, however, plateaus after *K* = 1, and every later adjacent MSE contrast has a bootstrap interval spanning zero. Residual energy is therefore the primary mechanistic evidence, and these results assign no independent biological meaning to any individual codebook.

### Paired token and prefill efficiency

The token audit pairs text and RVQ views of the same cells under the target checkpoint’s chat template, covering all 5,065 canonical prompts in the declared CTA, TI, DSA, and PCA scopes; every RVQ prompt is shorter than its paired text prompt.

These lengths also fix how many cells each modality carries under a fixed context budget. For both TI and DSA, budgets of 4,096, 8,192, and 16,384 tokens admit a median of at most 8, 16, and 32 cells through the text arm, whereas the RVQ arm fits the full *N* = 64 prompt at every one of them; the audit does not extrapolate beyond the observed *N* = 64 grid. At *N* = 64 the population strata use 96.1%– 96.5% fewer input tokens and show 38.51–41.02 × lower paired prefill latency. These measurements establish compactness under the specified tokenizer, prompts, checkpoint, hardware, and serving configuration; they establish neither accuracy parity between modalities nor the same latency ratio under batching or concurrency.

### Cross-method tokenizer comparison

The configuration study of §6.1 selects a shape within the residual-quantization family. Table 15 instead compares quantizer families on a balanced held-out set of 23,866 cells, grouped by information budget so that representation rates stay visible. The deployed 8 × 32 residual-VQ checkpoint is evaluated at its released step 25,500; Product Quantization (PQ), which partitions the latent and quantizes each block independently, and the matched CellTok-style reimplementation are evaluated at step 15,000, where their evaluation losses had plateaued.

§2 cites the utilization contrast between the matched CellTok-style single-codebook reimplementation (4.03%) and the deployed 8 × 32 residual cascade (85.94%). The grouped rows make its terms explicit: the two sit at different bit rates (256 versus 40) and different checkpoint steps, and “matched” refers to the shared reimplementation and held-out evaluation pipeline rather than to an equal information rate. Within the 256-bit group, PQ leads utilization and expression correlations while the CellTok-style reimplementation leads reconstruction MSE and clustering despite its low utilization; the higher-budget RVQ rows use 17.99%–22.36% of their enlarged vocabularies.

## G Systems Not Included in the Cross-Model Comparison

This subsection records, for the candidates that the single-cell community might expect, the per-system reason each is absent from Table 5. The inclusion criterion of §4.5 is a released native workflow supplying a task-compatible output contract; the omissions below do *not* share a single cause, and we separate them so that none is misread as a generic “protocol-incompatible” verdict.

### Protocol-bridge cases (scGPT, Geneformer)

**scGPT** [5] and **Geneformer** [6] produce cell embeddings rather than labels. A categorical prediction therefore requires either fine-tuning a classification head on the target cell types or attaching a nearest-prototype head over cell-type-name embeddings from an external text encoder; both routes introduce a head-choice or text-encoder confound that native-workflow systems do not carry. **LangCell** [14] ships its own published bridge (PubMedBERT + Geneformer), but its nearest-prototype scoring emits a similarity ranking over a fixed label vocabulary rather than the free-form output the deterministic graders parse.

### Task-adaptation case (CellFM)

**CellFM** [25] requires fine-tuning the 80M variant on the target labels for cell-type annotation; the 800M variant supports zero-shot inference only for gene-level tasks (gene function, GO multiclass), not cell-type labels [25]. The native release therefore does not provide an end-to-end cell-type label under the same conditions as the primary baselines.

### Capacity-scaled sibling (C2S-Scale-Gemma2-27B)

**C2S-Scale-Gemma2-27B** [13] satisfies the output-contract criterion: it produces labels under the same text-generation protocol as its 2B sibling and has been publicly available since October 2025. Its absence from Table 5 is therefore a scope decision rather than a protocol one. We report the parameter-matched member of the family, C2S-Scale-Gemma-2-2B (2.0B); at 27B parameters the larger variant is 6.8 × the size of our 4.0B backbone and would widen a scale range that the table already leaves uncontrolled.

## H CAPSTONE Benchmark Positioning

This appendix clarifies how CAPSTONE’s model-facing interface relates to existing single-cell evaluation resources.

### Benchmark positioning

Existing resources expose complementary interfaces. scBench gives agents direct access to .h5ad expression data for general scRNA-seq analysis [46]; SOAR evaluates cell-type annotation from marker lists [43]; and SC-Arena represents cells as ranked-gene sentences for multitask language-model evaluation [44]. CellVerse organizes cell, drug, and perturbation questions as multi-omics QA text [45], whereas VCWorld combines a knowledge graph with retrieved samples for perturbation reasoning [42].

CAPSTONE instead exposes expression-traceable RVQ tokens and, where required, a matched text view of the same biological instance across CTA, TI, DSA, and PCA. This distinction concerns the model-facing input contract rather than overall benchmark quality: the resources above serve different analytical scopes, and none is designed around paired text and RVQ views of one instance.

A separate line of perturbation-reasoning benchmarks evaluates capabilities outside CAPSTONE’s annotation and population scope, including differential-expression and direction prediction [54], regulation QA derived from LINCS [55], phenotypic-screen ranking [56, 57], and contrastive evidence for perturbation claims [58].

